# Tumor microenvironment immunomodulation by nanoformulated TLR 7/8 agonist and PI3k delta inhibitor enhances therapeutic benefits of radiotherapy

**DOI:** 10.1101/2024.03.09.584084

**Authors:** Mostafa Yazdimamaghani, Oleg V. Kolupaev, Chaemin Lim, Duhyeong Hwang, Sonia J. Laurie, Charles M. Perou, Alexander V. Kabanov, Jonathan S. Serody

**Affiliations:** Lineberger Comprehensive Cancer Center, University of North Carolina at Chapel Hill, Chapel Hill, NC, USA; Center for Nanotechnology in Drug Delivery and Division of Pharmacoengineering and Molecular Pharmaceutics, Eshelman School of Pharmacy, University of North Carolina at Chapel Hill, Chapel Hill, NC, USA; Department of Medicine, University of North Carolina School of Medicine, Chapel Hill, NC, USA; Department of Microbiology and Immunology, University of North Carolina, Chapel Hill, NC, USA

**Keywords:** Cancer Immunotherapy, Immunomodulation, Tumor microenvironment, Drug delivery, Nanomedicine, Radioimmunotherapy

## Abstract

Infiltration of immunosuppressive cells into the breast tumor microenvironment (TME) is associated with suppressed effector T cell (Teff) responses, accelerated tumor growth, and poor clinical outcomes. Previous studies from our group and others identified infiltration of immunosuppressive myeloid-derived suppressor cells (MDSCs) and regulatory T cells (Tregs) as critical contributors to immune dysfunction in the orthotopic triple-negative breast cancer (TNBC) tumor model limiting the efficacy of adoptive cellular therapy. However, approaches to target these cells specifically in the TME are currently lacking. To overcome this barrier, polymeric micelles nanoparticles (PMNPs) were used for co-delivery of small molecule drugs activating Toll-like receptors 7 and 8 (TLR7/8) and inhibiting PI3K delta. The immunomodulation of the TME by TLR7/8 agonist and PI3K inhibitor altered macrophage polarization, reduced MDSC accumulation and selectively decreased tissue-resident Tregs in the TME, while enhancing the T and B cell adaptive immune response. PMNPs significantly enhanced the anti-tumor activity of local radiation therapy (RT) in mice bearing orthotopic TNBC tumors compared to RT alone. Taken together, these data demonstrate that RT combined with a nanoformulated immunostimulant restructured the TME and has promising potential for future translation combined with RT for patients with TNBC.

## Introduction

TNBC is defined by the lack of estrogen and progesterone receptors and the absence of overexpression of human epidermal growth factor receptor 2. TNBC accounts for 10-15% of all breast cancers and is characterized by high tumor cell proliferation, abundant heterogeneity, frequent incidence of lung and brain metastases, repeated disease recurrence, poor prognoses, and is considered the most aggressive subtype of breast cancer [1–3]. In the absence of effective targetable biomarkers, chemotherapy in combination with surgery and radiation therapy (RT) continues to be the mainstay of standard of care for all patients with TNBC [4, 5]. Immune-enriched tumors have better outcomes with chemotherapy and patients with high expression of programmed death ligand 1 (PD-L1) benefited from immune checkpoint inhibitor (ICI) monoclonal antibody (mAb) therapy [4, 6]. Currently, the administration of anti-programmed death receptor 1 (PD-1) mAb in combination with cytotoxic chemotherapy is the backbone of neoadjuvant and metastatic therapy for patients with TNBC as single agent ICI lacks clinical efficacy for this group of patients [7]. The lack of effective immunotherapy approaches for patients with TNBC highlights the need for new strategies, including combinatorial treatments for targeting both tumor cells and non-cancer cells within the TME to improve TNBC therapeutic outcomes [4, 8].

Considering the molecular features and infiltrating immune cells, most TNBC tumors predominantly fall under two specific subtypes: basal-like and claudin-low [9]. Claudin-low breast tumors are associated with poor prognosis, high-grade large tumor size, and younger age of onset [10]. The claudin-low TNBCs are distinguished from other intrinsic molecular subtypes of breast cancer by low expression of cell-cell adhesion claudin-3, -4, and -7 genes, as well as genes encoding proteins involved in formation of tight junctions. Claudin-low tumors are enriched for genes linked with tumor cell proliferation, epithelial-to-mesenchymal transition (EMT), and stem cell-like characteristics, all of which are associated with poor patient outcomes [11, 12]. Thus it’s imperative to develop novel therapies to enhance the therapeutic options for patients with the claudin-low subtype. In mice, PD-1 and cytotoxic T lymphocyte-associated protein 4 (CTLA-4) targeting treatments are unsuccessful in suppressing tumor growth even though these tumors are enriched in adaptive immune cells [13].

Extensive evidence from translational studies indicates that immunosuppressive cell infiltration into the TME inhibits an efficient antitumor immune response [13]. Previously, we have shown that a large proportion of tumor-infiltrating lymphocytes in the claudin-low subtype of TNBC are CD4^+^Foxp3^+^ Tregs and tumor-associated macrophages (TAMs), leading to an immunosuppressive TME phenotype. Tregs in the TME interact with tumor cells, extracellular matrix components, stromal cells, and tumor-infiltrating cells. Additionally, they can secrete immunosuppressive mediators such as interleukin 10 (IL-10), IL-35, transforming growth factor β (TGF-β), and prostaglandin E2 (PGE2) [14]. The enrichment of Tregs in the TME diminishes cytotoxic effector T cell activity, contributing to tumor immune evasion by upregulating checkpoint inhibitor genes [15] and modulating transcription factors in TAMs [16] leading to chemotherapy resistance and early recurrence of clinical disease. One approach for remodeling immunosuppressive TME is centered on eliminating Tregs as their depletion in combination with anti-PD-1 and anti-CTLA-4 checkpoint inhibition increased cytokine production by CD8^+^ T cells, decreased tumor growth, and improved survival [13]. However, Treg-specific depletion in the TME has been quite challenging as these cells lack specific cell surface markers that can be used to target them for elimination and many current approaches are not specific to allow targeting of tumor-infiltrating Tregs.

Tregs are not the only cells that contribute to the formation of the immunosuppressive TME. Myeloid-derived suppressor cells (MDSCs) and TAMs promote tumorigenesis, immune surveillance evasion, angiogenesis, and metastasis [17]. The prevalence of TAMs in claudin-low subtype TNBC renders them a compelling focus for remodeling the TME via the polarization of immunosuppressive M2-like macrophages to an immunostimulatory M1-like fate. Previously, we reported that the modulation of the TME through STING agonists DMXAA or cGAMP shifted the balance of suppressive myeloid cells to immunostimulatory cells and enhanced CAR-T cell persistence, achieving tumor regression [18]. Along with other research groups, we demonstrated impeding macrophage recruitment, proliferation, and survival with a colony-stimulating factor 1 receptor (CSF1R) inhibitor, combined with chemotherapy agents such as paclitaxel or cyclophosphamide, resulted in repolarization of the immunosuppressive TME, enhance adaptive T cell immunity, and long-term posttreatment tumor regression in TNBC models [19–21]. These findings indicate that patients may significantly benefit from combining primary standard of care with immunomodulators to balance immunosuppressive and immune-activating factors in the TNBC TME.

The well-defined safety profile, widespread clinical availability, and potential immunostimulatory effect of radiation therapy (RT) make it a primary candidate for combination therapy with immunotherapy. Radiotherapy alters both cancer and immune cells in the TME and, depending on dose, regimen, and target volume, can induce immunostimulatory or immunosuppressive effects. In addition to decreasing the tumor burden and improving local extravasation and penetration of nanoparticles into TME [22], RT can synergize with immunotherapy, inducing infiltration of immune cells recruited by secreted inflammatory mediators from irradiated cells. However, RT is a double-edged sword and has been associated with an increase in immunosuppressive cells in the TME, inducing radioresistance through the induction of cellular senescence as senescent cancer cells demonstrate pro-tumorigenic properties via the secretion of immunosuppressive cytokines, resulting in the accumulation of immunosuppressive M2-like TAMs, MDSCs, Tregs, and induction of T cell exhaustion. [23, 24]

While multiple approaches have been attempted to modulate the immunosuppressive TME, many target populations of immune cells systemically, not solely in the TME or tumor draining lymph node TDLN, or target only a single population of immunosuppressive cells. Here, we report the development of an approach to broadly alter the immunosuppressive TME found in breast tumors. PMNPs co-loaded with TLR 7/8 agonist Resiquimod and the PI3K delta inhibitor idelalisib markedly potentiated RT, which was associated with significant changes in CD4^+^ and CD8^+^ T cells as well as B cells and myeloid cells. As this approach uses FDA-approved medications, it could be rapidly translated to enhance RT treatment of patients with claudin-low tumors TNBC. Finally, we provide a sophisticated immune assessment of the effects of different proteins loaded onto PMNPs using high dimensional spectral flow cytometry of the TME and TDLN.

## Results and Discussion

### Co-delivery of TLR7/8 agonist and PI3K inhibitor with nanoparticles for TME immunomodulation

To develop approaches to comprehensively alter the immunosuppressive TME, we used the murine syngeneic transplant T11-APOBEC3C (T11-AP) tumor model as previously published by our group [25]. We employed combination RT and nano-formulated immunotherapy as a treatment strategy for claudin-low TNBC. For this, R848 or resiquimod (Res), a potent imidazoquinoline Toll-like receptor (TLR) 7/8 agonist and idelalisib (ID), an inhibitor of phosphatidylinositol 3-kinase (PI3K) p110δ, were used. Polymeric micelle formulations of Res and ID were prepared using a thin film method (**Fig. 1a**). As both small-molecule drugs exhibit minimal solubility in water, we incorporated them in poly(2-oxazoline) block copolymer micelle nanoparticles (PMNPs) [26]. We hypothesized that NPs delivering immunostimulatory compounds could be efficacious in the claudin-low TNBC subtype, as Tregs and immunosuppressive myeloid cells make up the majority of the tumor immunosuppressive composition (**Fig. 1b**).

We incorporated Res and ID separately in the POx micelles with high loading efficacy (LE) of 92.4% and 99.3%, respectively. The high loading capacity (LC) of 42.4% for Res and 44.2% for ID in single-loaded micelle potentiates delivering a high drug dose with a small POx polymer content. The Res and ID combination micelles (Res:ID) were co-loaded in nanoparticles at a 1/4 (w/w) drug ratio with 98.7% and 94.9% LE, indicating little loss of the drugs upon loading. A substantial amount of each drug was incorporated in co-loaded micelles, with combined LC of 51.4%: 13.6% of Res, and 37.8% LC for ID (**Fig. 1c**). To evaluate the size, size distribution, and morphology of particles of the single loaded micelles and co-loaded nanoparticles, we employed dynamic light scattering (DLS) and transmission electron microscopy (TEM). The resulting Res micelle, ID micelle, and Res:ID micelle were spherical (**Fig. 1d**) with particle sizes of 27.6 nm, 33.8 nm, and 32.9 nm, respectively, with PDI of 0.21, 0.14, and 0.24, indicating monodispersity of prepared spherical nanoparticles (**Fig.1c**).

To assess the potential stability of PMNPs loaded with Res, we measured the particle size at room temperature of micelle solutions stored at 4°C for up to 18 days by DLS. The single drug Res micelles exhibited increased particle size and polydispersity index (PDI), accompanied by visual drug precipitation on day 1, demonstrating instability of PMNPs with Res. Conversely, the co-loaded Res:ID micelles were stable during the entire observation period, retaining small size and PDI with no observable precipitation (**Fig. 1e**). Given the robust stability of ID micelles of at least 18 days and ID to Res drug ratio of 4:1, we hypothesized that the ID interactions with POx polymer increased the colloidal stability of co-loaded Res:ID micelles. Next, we evaluated drug release profiles under sink conditions and observed similar release profiles for Res and ID from single-loaded and co-loaded micelles, with 80% of drug released at 10 h (**Fig. 1f**). Thus, we generated PMNPs with enhanced stability and similar release profile by co-loading Res with ID.

**Fig. 1.**
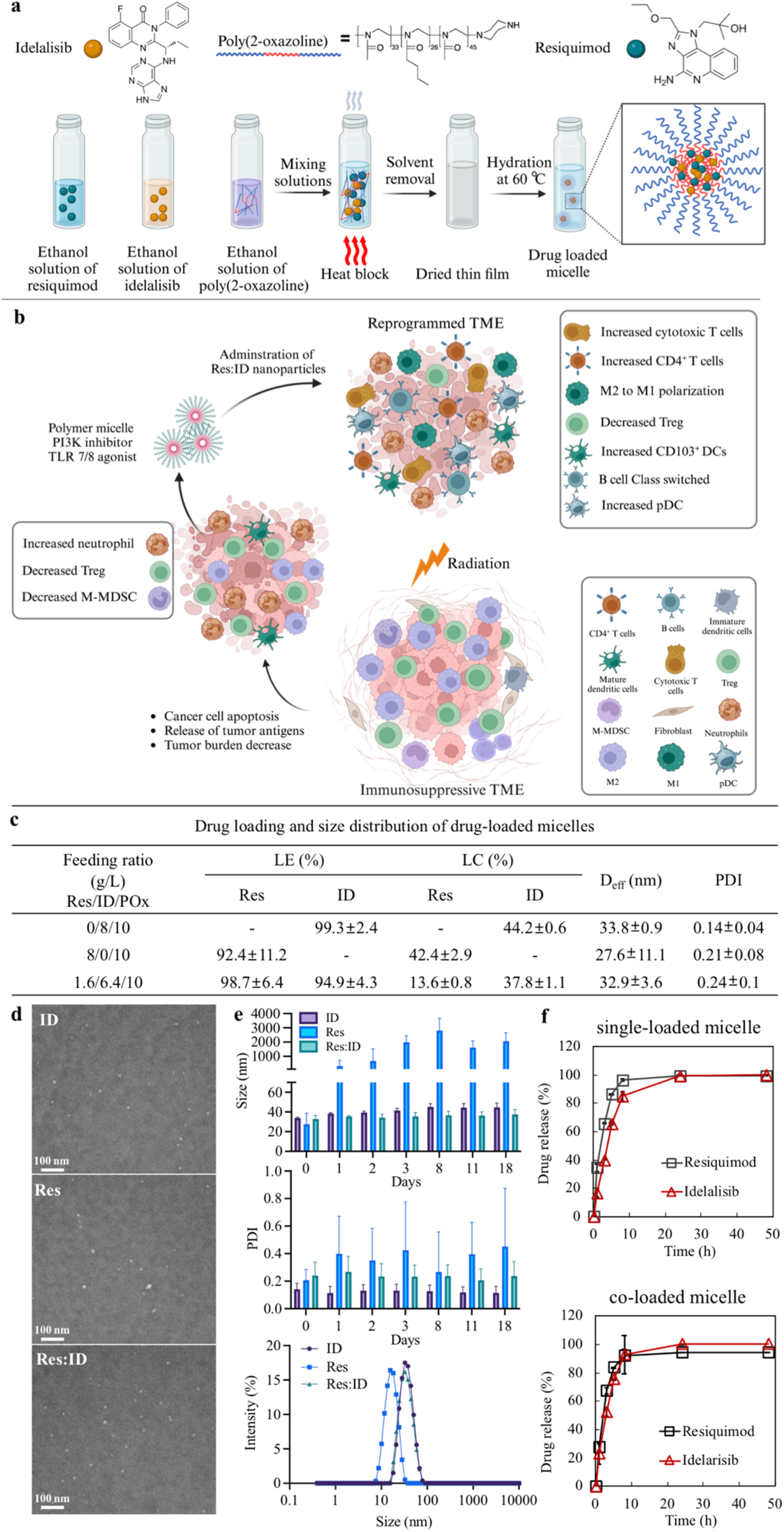
Synthesis and characterization of drug-loaded polymer micelle nanoformulation. **a)** Schematic of drug-loaded polymer micellar preparation and **b)** proposed mechanism of tumor microenvironment immunomodulation by TLR7/8 agonist and PI3k inhibitor for enhanced cancer immunotherapy. **c)** Drug loading (% loading efficiency (LE) = M_drug_ / M_drug added_ × 100 and % loading capacity (LC) = M_drug_ / (M_drug_ + M_excipient_) × 100) and size distribution (D_eff_ and polydispersity index (PDI)) of micelles. **d)** Drug-loaded micelles stability evaluation by dynamic light scattering (DLS) over time by monitoring particle size and PDI change. **e)** Representative transmission electron microscopy (TEM) images and **f)** drug release profiles from single drug micelles and co-loaded micelles.

### Immune responses to nanoformulated TLR7/8 agonist and PI3K inhibitor

We next assessed the efficacy of these nanoformulated drugs in our T11-AP model. For this assessment, a PBS control group or recipients of ID (80mg/kg) PMNPs, Res (20mg/kg) PMNPs and Res:ID (20:80 mg/kg) PMNPs, were evaluated for tumor growth and survival (**Fig. 2a-c**). While none of the treatments showed significant suppression of tumor growth (**Fig. 2b-c**), co-loaded PMNPs delayed tumor growth by up to 10 days and decreased tumor burden compared to the control groups (**Fig. 2c**). Flow cytometric analysis of homogenized tumors revealed significant changes in the TME. The ID-containing treatment modestly decreased the frequency of CD25^+^ Foxp3^+^ Treg (**Fig. 2e**), while co-loaded PMNPs significantly promoted infiltration of activated CD8^+^ T cells and CD8^+^ Tem in the TME (**Fig. 2e, f**). Res-containing treatments increased the frequency of plasmacytoid dendritic cells (pDCs) in the tumor (**Fig. 2g**). We observed TLR7/8 agonist treatment significantly increased M1 macrophages while slightly decreasing the M2 population (**Fig. 2i, j**), which correlated with augmented iNOS expression and reduced CD206 expression on macrophages in TME (**Fig. 2k**).

While Res treatment enhanced M1 macrophages it was also associated with a significant increase in CD11b^+^Ly6C^+^Ly6G^-^iNOS^+^CD103^-^ monocytic MDSCs (M-MDSC) in the TME (**Fig. 2h**) versus granulocytic MDSCs (G-MDSCs, CD11b^+^Ly6C^low^Ly6G^+^CD101^-^CD84^+^, distinguished from granulocytes by CD101 expression). This increase in M-MDSCs was found with single and dual loaded PMNPs and confirmed previous data indicating that TLR agonists can enhance MDSC numbers and function. [27–30]

**Fig. 2.**
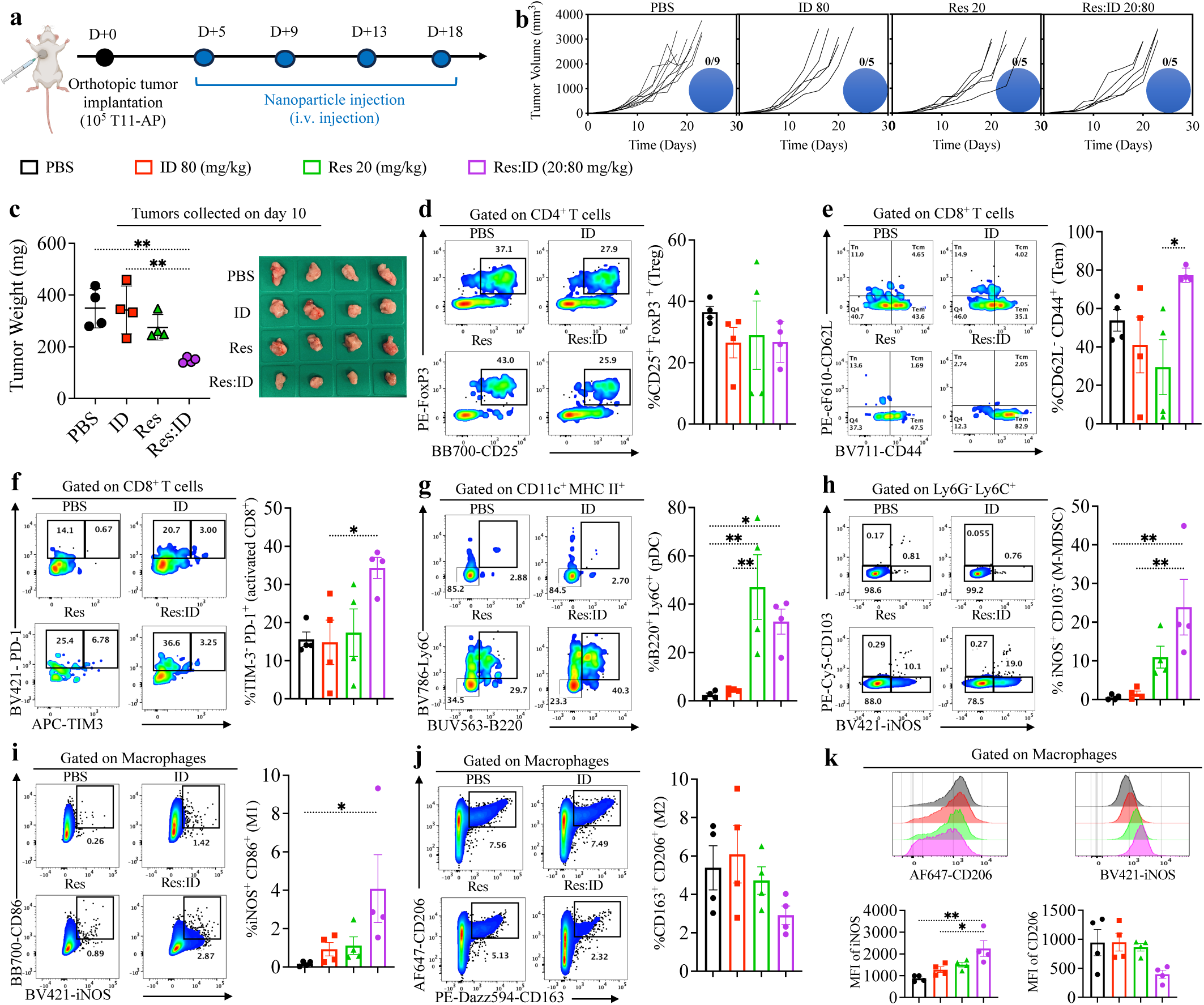
In vivo antitumor therapeutic effect and immune responses to i.v. administration of ID80, Res20, and Res:ID20:80 in orthotopic T11-Apobec tumor model. **a)** Schedule of formulation treatment. T11-Apobec tumor-bearing Balb/c mice were treated intravenously with single-loaded or co-loaded nanoparticles, on days 5, 9,13, and 18, and **b)** individual tumour growth and **c)** The tumor weights and tumor images with different treatments. Tumor tissues were harvested on day 10 after 2 times i.v. administration of nanoformulated drugs. **d-k)** Representative flow cytometry dot plots and corresponding bar graphs for immune cell response. All data are presented as the mean ± SEM with n=4. **d)** The TME was analyzed on day 10 by flow cytometry for the frequency of CD4^+^FoxP3^+^ Treg cells, **e)** effector memory CD8^+^ T cells (CD62L^-^CD44^+^), **f)** PD-1^+^ Tim3^-^ activated CD8^+^ T cells, **g)** plasmacytoid dendritic cells (pDCs), **h)** CD103^-^iNOS^+^ M-MDSC, **i)** CD86^+^iNOS^+^ M1-like macrophages, **j)** CD206^+^CD163^+^ M2-like macrophages, **k)** iNOS and CD206 expression on macrophages within the TME. Ordinary one-way ANOVA followed by a Tukey’s multiple-comparisons test was applied for statistical analyses. *P* values: NS, not significant, \**P* < 0.05, \*\**P* < 0.01, \*\*\**P* < 0.001, \*\*\*\**P* < 0.0001).

### Radiotherapy in combination with Res:ID20:80 eliminates established TBNC tumors

Idelalisib is a selective inhibitor of the class I PI3K p110δ isoform (PI3Kδ). Although the α and β isoforms are expressed widely, the δ and γ isoforms are selectively expressed on leukocytes. Idelalisib received accelerated approval from the FDA for the treatment of several B-cell malignancies but was later withdrawn from the market, in part, due to a challenging safety profile that included hepatoxicity and serious pneumonitis. To address these issues, we assessed drug toxicity in various concentrations, administration routes, and in combination with RT and TLR7/8 agonist to ensure that the proposed doses and regimens were safe (**Supplementary Fig. 2S-4S**). Interestingly, the increased hepatotoxicity found using PMNPs loaded with ID was not observed in the co-loaded PMNPs with ID and Res. Thus, by co-loading PMNPs, the stability of Res in the NP was enhanced while the toxicity of ID was decreased.

Given the alteration of the TME with the PMNPs co-loaded with Res and ID, we were interested in evaluating whether this approach synergized with RT in mice with T11-AP tumors. We hypothesized by encapsulating PI3Kδ in nanoparticles, we could leverage the known enhanced permeability and retention (EPR) effect to target TME, thereby curtailing off-target toxicity, selectively decreasing the number and function of Tregs in tumors, and limiting the frequency of drug administration. Strikingly, RT significantly extended the median survival from 20 days to 37 days in the RT compared to in PBS group, respectively (**Fig. 3a**). While ID given at 20 and 80 mg/kg combined with RT (RT+ID) delayed tumor growth and extended median survival, ID monotherapy showed no benefit, and recipients of RT treatment alone exhibited tumor growth and median survival similar to RT+ID (**Fig. 3b-c**) with no complete responders. As recent studies have demonstrated the efficacy of TLR agonists to enhance anti-tumor immunity [31, 32], we next treated mice with Res alone or in combination with RT at low (5 mg/kg) and high concentrations (20 mg/kg). Though Res monotherapy modestly delayed tumor growth, animals treated with the combination of RT and the high-dose Res (RT+Res20) displayed significant tumor inhibition with eradication of tumors in a minority of treated mice (**Fig. 3b, d**). Although treatment with the low-dose Res in combination with RT (RT+Res5) suppressed tumor growth significantly, no animals exhibited complete tumor elimination and all mice eventually succumbed due to tumor growth (**Fig. 3b, d**). However, there was a significant enhancement of the anti-tumor effect of RT in mice treated with PMNPs co-loaded with both drugs at high concentrations (RT+Res:ID 20:80). Mice with T11-AP tumors receiving RT+Res:ID 20:80 had a significant improvement in survival with eradication of tumor in 50% of mice while the lower-dose combination (RT+Res:ID 5:20) had antitumor effects indistinguishable from RT alone (**Fig. 3b, e**). Interestingly, RT treatment combined with oral or i.v. administration of free drug formulations of Res and ID in a standard vehicle at high dose showed a comparable antitumor effect to RT monotherapy and was significantly inferior to the administration of Res/ID in PMNPs (**Fig. 3b, f**). Thus, the combination of high dose nanoformulated Res and ID in PMNPs given with RT was significantly better than the clinical routes of oral or i.p. administration of the free drugs with RT.

**Fig. 3.**
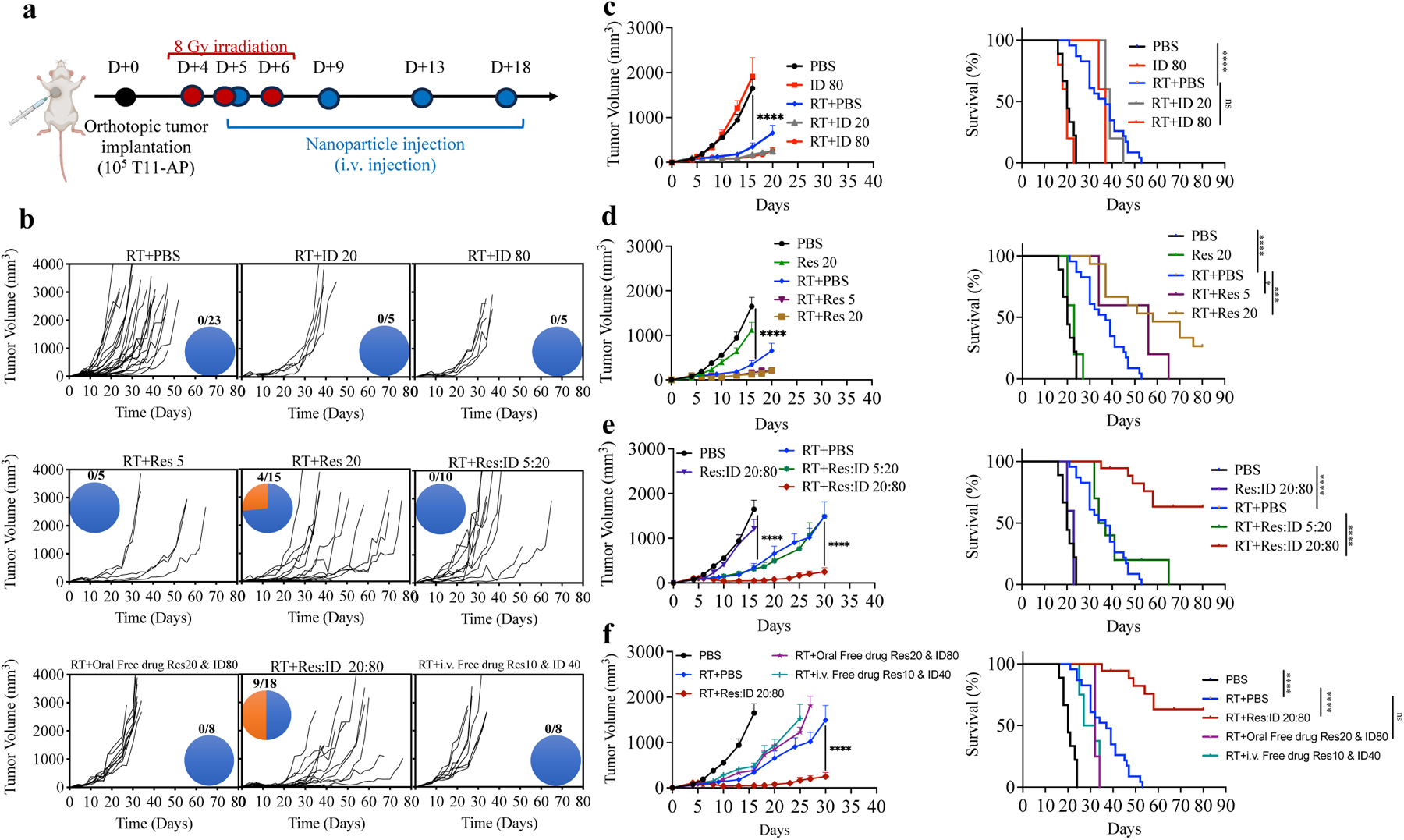
Co-delivery of TLR7/8 agonist and PI3K inhibitor within nanoparticles in combination with radiotherapy eliminates established TNBC tumors. **a)** Schedule of formulation treatment and RT dosing. T11-AP tumor-bearing Balb/c mice received targeted 8 Gy radiation on days 4, 5, and 6, following intravenous nanoformulated TLR7/8 agonist and PI3K inhibitor either as single-loaded or co-loaded drugs on days 5, 9,13, and 18. **b)** Individual tumor growth, number of animals per group, complete response rate, **c-f)** average tumor size and percentage of survival were monitored over time. All data are presented as the mean ± SEM. Statistical comparisons for tumor volume were analyzed using Mann-Whitney unpaired two-tailed t-test, and survival percentages were presented as Kaplan-Meier survival curves and analyzed via Log-rank (Mantel-Cox) test. *P*values: NS, not significant, \**P* < 0.05, \*\**P* < 0.01, \*\*\**P* < 0.001, \*\*\*\**P* < 0.0001).

### The Res:ID20:80 co-loaded NPs modulate lymphocytes and myeloid cell profile to improve radiotherapy

To evaluate how the Res and ID co-loaded PMNPs mediated their effect when combined with RT, cells from the TME and tumor-draining lymph node (TDLN) were assessed using spectral flow cytometry. We designed a 26-parameter spectral flow cytometry panel for immunophenotyping major immune cell subsets in the tumor and TDLN of T11-AP tumor-bearing mice treated with PBS, RT, RT+ID80, RT+Res20, and RT+Res:ID 20:80 (**Fig. 4**). The heatmap of the marker expression, abundance of lymphocyte clusters, and density plots of selected marker expression used for cluster identification and annotation are in **Fig. 4b, e and Supplementary Figures 5S-7S**, respectively. Subsequently, identified clusters were analyzed to define significant immune profile changes between treatments. Eight main populations of tumor infiltrating immune cells, including B cells, CD4^+^ T cells, CD4^+^ Tem, Tregs, CD8^+^ T cells, MHC II^Hi^ DCs, MHC II^low^ DCs, and CD103^+^ migratory DCs, were conserved among treatment groups. Recipients of Res:ID co-loaded PMNPs + RT showed significant changes in Treg and B cell phenotype. In addition to a modest reduction of Treg frequency, we observed decreased expression of CD103, a marker for the tissue-resident Tregs, compared to the RT group (**Fig 4c**). Mice receiving PMNPs with Res had increased expression of the activation marker PD-1 on B cells along with higher expression of CD138 on B cells and CXCR5 expression on CD4^+^ T cells, suggesting the augmented cross-talk between follicular helper T (Tfh) cells and antibody-secreting cells in response to treatment (**Fig. 4c**). In addition, we noted RT treatment induced an increase in frequency of MHC II^low^ DCs subpopulation in the TME compared to PBS controls. In the RT+Res treatment more CD103^+^ migratory DCs infiltrated the TME and while there were limited differences between RT and RT+ID, a higher frequency of MHC II^low^ DCs and a lower percentage of Tregs were present in RT+ID treatment (**Fig. 4c**).

We further explored the shifts in the immune profile of the TDLN in response to treatments. In total, we identified nine discrete clusters of lymphoid cells within the TDLN, corresponding to CD4^+^ Tems (CD62L^-^CD44^+^ effector memory T cells), CD4^+^ Tn (CD62L^+^CD44^-^ naive T cells), CD4^+^ Tcms (CD62L^+^CD44^+^ central memory T cells), CD4^+^ CD62L^-^ CD44^-^ T cells, CD8^+^ Tem, CD8^+^ Tn, Tregs, IgD^+^IgM^+^ B cells, and IgD^-^IgM^+^ B cells (**Fig. 4d, e**). Mice treated with PMNPs co-loaded with Res and ID in combination with RT exhibited higher frequency of CD8^+^ Tn, CD8^+^ Tems, and CD4^+^ Tems in their TDLN compared to PBS control and RT alone groups, indicating that TLR7/8 and PI3K treatment activated the adaptive anti-tumor immune response (**Fig. 4f**). Interestingly, unlike in the TME, Treg frequency in the TDLN increased with TLR7/8 treatment, though this was offset by increased infiltration of CD8^+^ T cells, leading to no change in the CD8^+^ /Treg ratio compared to control mice (**Fig. 4f**). Moreover, concomitant with a decrease in the frequency of IgD^+^IgM^+^ B cells, the percentage of the IgD^-^IgM^+^ B subset increased for RT+Res:ID group. Similar to the TME, CXCR5 expression was increased on CD4^+^ T cells in the TDLN, with expression of PD-1 and CD138 on B cells increased in the RT+Res:ID group, indicating that combined TLR stimulation and PI3K delta inhibition induced B cell activation and differentiation. (**Fig. 4f**). These data suggest that part of the antitumor effect of Res occurs via activation of TLR7 and TLR8 expressed in B cells, concomitant with decreased Treg function in the TME mediated by ID, which maintained T cell anti-tumor activity [33]. In addition, increased CXCR5 expression on CD4^+^ T cells, and PD-1 and CD138 on B cells could indicate enhanced interactions between Tfh cells and antigen-secreting cells in TDLN of mice treated with co-loaded NPs.

**Fig. 4.**
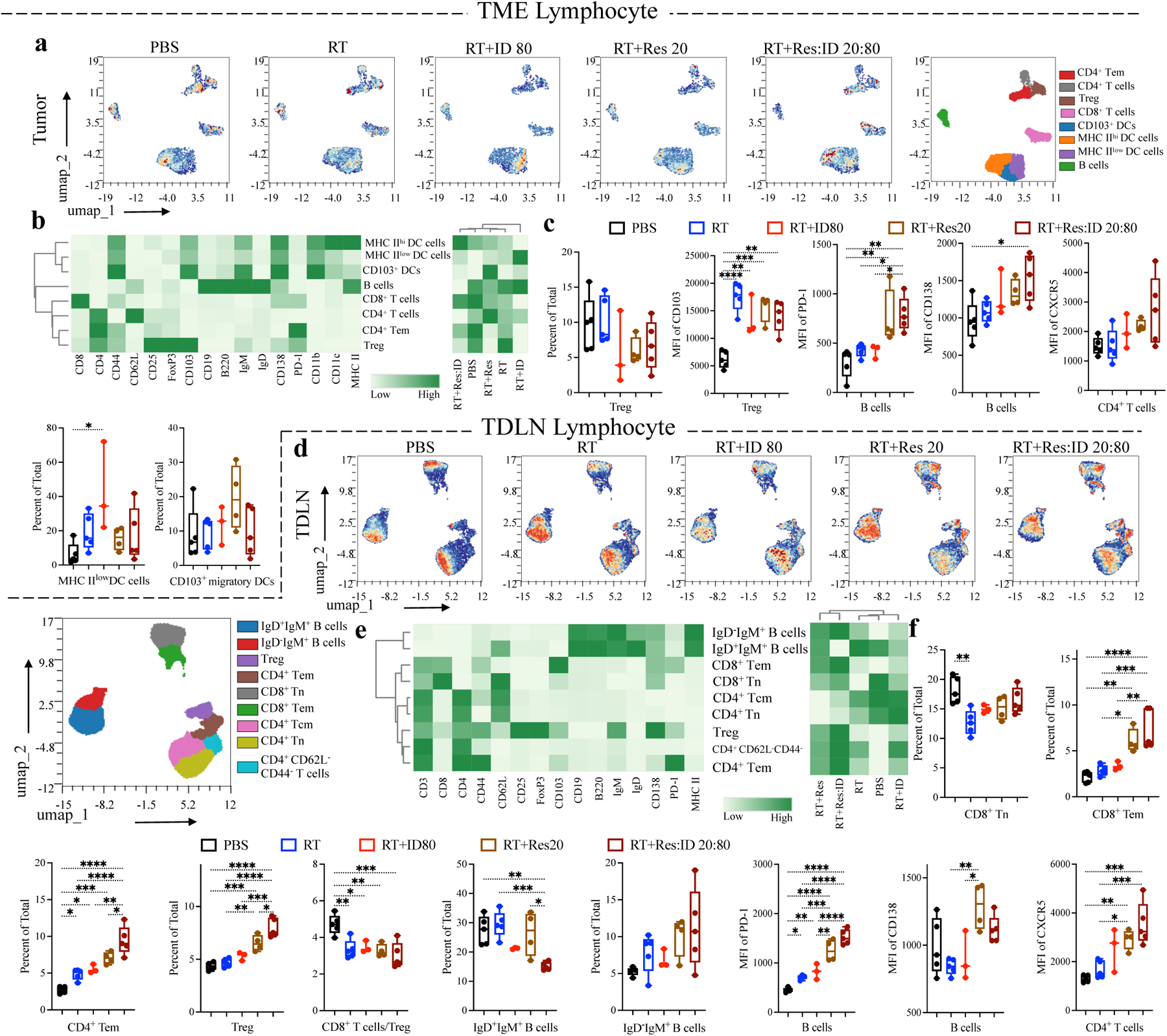
Characteristics of lymphoid cells landscape in response to the combination of radiotherapy and TLR7/8 and PI3K immunomodulators. High dimensional spectral flow cytometry analysis was conducted for an in-depth investigation of the immune landscape profile across different treatments in TME and TDLN. T11-AP tumor-bearing Balb/c mice received targeted 8 Gy radiation on days 4, 5, and 6, following systemic i.v. administration of nanoformulated TLR7/8 agonist and PI3K inhibitor on days 5, 9,13, and 18, and tumor and TDLN harvested on day 30 for spectral flow cytometry analysis. Control (n=5), RT (n=5), RT in combination with ID80 (n=3), RT in combination with Res20 (n=4), and RT in combination with Res:ID 20:80 (n=5) were analyzed to study the effect of radiation on lymphoid cells and nanoformulated drugs were compared with control and RT treatment. **a,d)** Uniform manifold approximation and projection (UMAP) density plots for a total of 30,000 lymphocytes in the tumor and 123,000 lymphocytes in TDLN were examined by spectral flow cytometry and analyzed using OMIQ. Myeloid subsets were excluded to increase clarity. FlowSOM clusters were projected on a UMAP plot, and each cluster were identified by the lymphoid panel, UMAP scatterplot of relative expression of major lineages markers (Fig. 6-8S), and **b,e)** clustered heatmap of the lymphoid markers’ expression. Significant changes in identified FlowSOM clusters in between treatments were recognized from abundance heat maps. **c,f)** Frequency of total for subpopulations of T cells and B cells, and expression level PD-1, and CD138 on B cells and CXCR5 on CD4^+^ T cells were compared between all treatments. All data are presented with ordinary one-way ANOVA followed by a Tukey’s multiple-comparisons test for statistical analyses. *P*values: NS, not significant, \**P* < 0.05, \*\**P* < 0.01, \*\*\**P* < 0.001, \*\*\*\**P* < 0.0001).

To further explore the effect of treatment on immune dynamics in the myeloid component of the TME, we utilized the spectral flow cytometry UMAP dimension reduction technique and FlowSOM clustering to investigate the relationship of myeloid cells to each treatment performed (**Fig. 5 and Supplementary Fig. 8S**). Animals exposed to either RT+Res20 or RT+Res:ID 20:80 treatments had a significantly higher frequency of neutrophils in the TME, indicating that the Res-containing PMNPs act synergistically with RT to enhance neutrophil frequency compared to control and RT alone. We also observed a significant decrease in the frequency of M-MDSCs in the TME (**Fig. 5c**) after RT, which offset the enhanced frequency of M-MDSCs observed in Fig. 2h after TLR7/8 stimulation. While the decrease in the frequency of M-MDSCs was observed to be less pronounced after TLR7/8 stimulation, the combination of radiotherapy with a 20:80 ratio of Res to ID (RT+Res:ID) showed a significantly lower number of M-MDSCs in the TME. We noted a similar effect on the pDC subset (**Fig. 5c**).

**Fig. 5.**
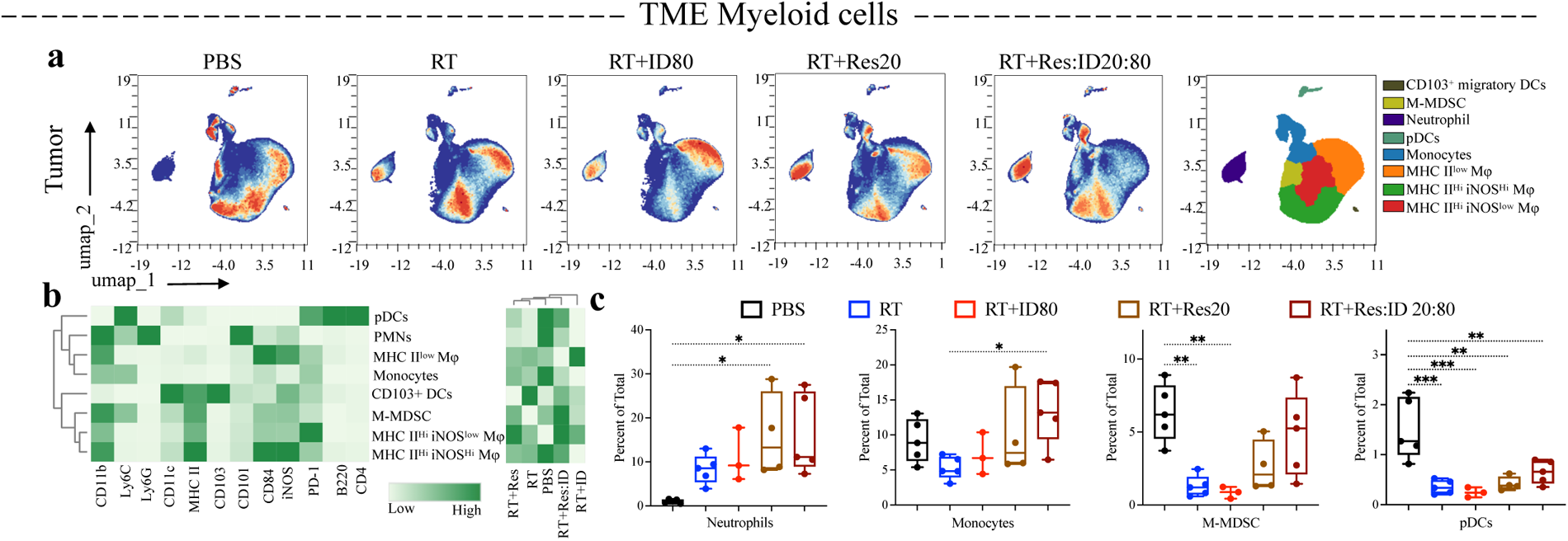
Characteristics of myeloid cells landscape in response to the combination of radiotherapy and TLR7/8 and PI3K immunomodulators. High dimensional spectral flow cytometry analysis was conducted for an in-depth investigation of the immune landscape profile across different treatments in TME. T11-AP tumor-bearing Balb/c mice received targeted 8 Gy radiation on days 4, 5, and 6, following systemic i.v. administration of nanoformulated TLR7/8 agonist and PI3K inhibitor on days 5, 9,13, and 18, and tumor were harvested on day 30 for spectral flow cytometry analysis. Control (n=5), RT alone (n=5), RT in combination with ID80 (n=3), RT in combination with Res20 (n=4), and RT in combination with Res:ID 20:80 (n=5) were analyzed to study the effect of radiation on myeloid cells and nano-formulated drugs were compared with control and RT treatment. **a)** UMAP density plots for a total of 1.2 × 10^6^ myeloid cells in the tumor were examined by spectral flow cytometry and analyzed by OMIQ. Lymphocyte subsets were excluded to increase clarity. FlowSOM clusters were projected on a UMAP plot, and each cluster were identified by the myeloid panel, UMAP scatterplot of relative expression of major lineages markers (Fig. 9S), and **b)** clustered heatmap of the myeloid cells markers’ expression. Significant changes in identified FlowSOM clusters in between treatments were recognized from abundance heat maps. **c)** Frequency of total subpopulations of PMNs, macrophages, monocytes, DCs, and M-MDSC were compared between all treatments.

Taken together the high-level analysis revealed synergistic anti-tumoral effects of PI3K inhibitor and TLR7/8 agonist delivery via co-loaded PMNPs post-radiation. We next examined populations of cells in the TME that were affected by the treatment by analyzing tumor weight-normalized absolute counts of these cells. Specifically, we observed a dramatic 2-3-fold increase in both total T cells, CD4^+^ and CD8^+^ T cells only in the Res:ID PMNP group. Strikingly these changes were primarily driven by the increases in infiltrating CD4^+^ and CD8^+^ T cells with an effector memory (CD44^+^CD62L^-^) phenotype (**Fig 6b**). Additionally, the number of activated CD8^+^ T cells that expressed PD-1 but lacked expression of TIM-3, a coinhibitory molecule found on exhausted T cells, was also significantly higher in Res:ID PMNP-treated group compared to all other conditions (**Fig. 6b**). Moreover, the greatest decrease in the ratio of Treg before and after RT treatment for each group was observed in mice receiving RT with Res:ID PMNPs (**Fig. 6b**). The same effect was noted in the ratio of Tregs to CD8^+^ T cells. These data demonstrate that dual Res:ID PMNPs in combination with RT specifically decrease immunosuppressive Treg cells while enhancing the numbers of CD4+ and CD8+ T cells. We also observed TLR-stimulation enhanced the number of B cells and, more specifically, class-switched B cells in the TME, suggesting that co-loaded NPs may promote antibody-secreting cells with the potential to target tumor-associated antigens (**Fig. 6c**). The synergistic effect of the Res:ID combination with RT revealed successful reduction of the ratio of M-MDSCs before and after RT for each treatment (**Fig. 6d**) compared to treatments without RT. Finally, polarization of M2 macrophages to M1 type and increased infiltration of CD103^+^ migratory DCs were observed in mice receiving co-delivery of RT with Res:ID PMNPs (**Fig. 6d**).

Given the significant changes in the TME mediated by RT with Res:ID PMNPs, we were interested in determining which of these changes was critical to the success of this therapy. Thus, we used depleting antibodies to assess tumor growth and survival of RT+Res:ID recipients compared to the same treatment in the animals deficient in either B cells, CD8^+^ or CD4^+^ T cells, or Gr1^+^ myeloid cells (**Fig. 6e**) [34]. Strikingly, the control of tumor growth was lost in RT+Res:ID-treated mice after depletion of either lymphoid (CD4^+^ and CD8^+^ T cells, B cells) or myeloid cells broadly, indicating an important role of these immune cells in the anti-tumor immune response. However, CD4^+^ T cells and myeloid lineage cells (Gr1^+^) were absolutely required as their depletion resulted in a dramatic reduction of survival. These data demonstrate that PMNPs loaded with Res:ID can activate the myeloid compartment to inhibit tumor growth without adversely affecting the B cell or T cell response critical for tumor control in response to RT. Taken together, these data support our hypothesis that RT+Res:ID 20:80 alters the TME to enhance the anti-tumor immune response. This correlated with changes to the TME that included changes in the polarization state of macrophages toward an M1 phenotype, increased number of neutrophils, and migrating DCs while enhancing the adaptive T cell and B cell immune response.

**Fig. 6.**
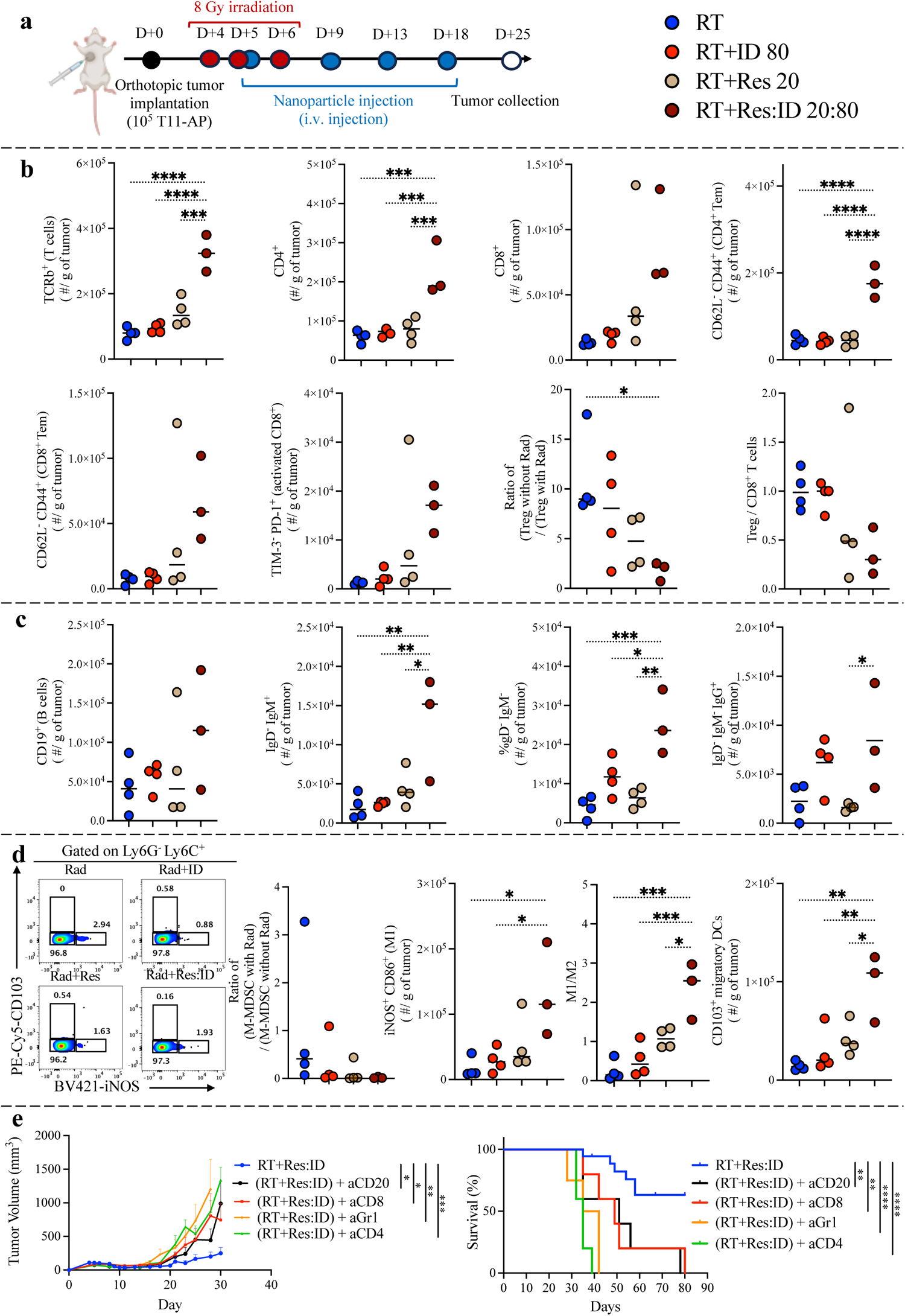
Treatment with Res:ID20:80 remodels the radiated TNBC tumor microenvironment. **a)** The Schedule for the treatment of Balb/c tumor-bearing mice. **b-d)** Flow cytometry analysis of the normalized number of T cells, CD4^+^ T cells, CD8^+^ T cells, effector memory CD4^+^ T cells, effector memory CD8^+^ T cells, Tim3^-^PD-1^+^ activated CD8^+^ T cells, Tregs, B cells, CD103^+^ migratory DCs, CD86^+^iNOS^+^ M1-like macrophages, ratio of M1-to M2 like macrophages, and representative flow cytometry dot plots CD206+CD163+ M2-like macrophages and CD103^-^iNOS^+^ M-MDSC within the TME. All results are presented as means ± SEM (n = 4). Statistical significance for flow data was calculated using ordinary one-way ANOVA followed by a Tukey’s multiple-comparisons test. **e)** Average tumor volume and survival plots for mice given RT and co-loaded NPs with Res and ID, in combination with either anti-CD8, anti-Gr1, anti-CD4, anti-CD19, or anti-CD20 antibodies. The RT group comprises 18 mice from 4 independent experiments, while the remaining groups have a sample size of n=4 or n=5. Statistical comparisons for tumor volume were analyzed using Mann-Whitney unpaired two-tailed t-test, and survival percentages were presented as Kaplan-Meier survival curves and analyzed via Log-rank (Mantel-Cox) test. *P*values: NS, not significant, \**P* < 0.05, \*\**P* < 0.01, \*\*\**P* < 0.001, \*\*\*\**P* < 0.0001).

## Conclusion

Understanding how to modulate the TME can enhance the efficacy of immunotherapy and clinical outcomes for patients with cancer. However, targeting the immunosuppressive TME has been quite challenging as there are very few approaches that globally alter the TME, and single-agent approaches lack tumor specificity. Administration of Res:ID in PMNPs abrogated the toxicity of ID and greatly enhanced the stability of Res in the PMNP. This allowed us to combine RT with Res:ID PMNPs to induce effective TME immunomodulation, which enhanced tumor regression and increased median survival and with over a 50% complete response rate. When compared to RT alone or RT with single-loaded PMNPs, administration of co-loaded PMNPs with Res:ID enhanced the number and function of effector T cells, and activated B cells in the TME and TDLN while greatly enhancing the M1/M2 ratio of macrophages. The radioimmunotherapy approach employed in this study effectively reduced the tumor burden, which was associated with modulating the TME through a dual mechanism. One aspect involved disrupting the TME by reducing the numbers of immunosuppressive cells such as Treg, MDSCs, and M2 macrophages. Simultaneously, the other component activated effector memory CD4^+^ and CD8^+^ T cells, enhancing the migration of Dendritic Cells (DCs), increasing M1 macrophages, and promoting cross-talk between Tfh cells and antibody-secreting B cells. Taken together, the work presented here highlights the role of immunomodulatory agents in improving the therapeutic benefits of RT and provides an understanding of immune cell modulation for efficient immunosuppressive TME remodeling as a promising therapeutic approach for treating immunosuppressed breast tumors.

## Materials and Methods

### Materials

Living cationic ring-opening polymerization was used to synthesize amphiphilic triblock copolymer of poly (MeOx_35_-b-BuOx_20_-b-MeOx_34_) (Mn = 8.6kDa, Mw/Mn = 1.15), as described previously.[19] Gel permeation chromatography (GPCmax VE-2001 system (Viscotek)) and ^1^H NMR spectroscopy (INOVA 400) were utilized to characterize the structural properties of POx. Resiquimod (R848) and idelalisib (CAL-101; GS-1101) were obtained from MedChemExpress (Monmouth Junction, NJ, USA). T11-AP cells were cultured in RPMI medium (11965–092 (Gibco)) supplemented with 1% penicillin-streptomycin, 1% L-Glutamine, 1% Sodium Pyruvate, 1% Non-Essential Amino Acids, 5% FBS, 1% 2-Mercaptoethanol, and puromycin. The list of antibodies used in flow cytometry and depletion studies is summarized in Table 1S.

### Formulation and characterization of polymeric micelles loaded with small molecule drugs

Fig. 1 presents a schematic of drug-loaded polymer micellar preparation and characterization of drug-loaded polymer micelle nanoformulation. As previously outlined, the thin-film hydration method produced POx micelle formulations containing drugs [19]. In summary, stock solutions of the POx polymer and drugs (Res and ID) were prepared in absolute ethanol, each at a concentration of 10 mg/mL. To obtain a definite drug-feeding ratio in POx micelles, predetermined ratios of stock solutions were mixed, and ethanol content was evaporated entirely under a stream of inert nitrogen gas. The prepared homogeneous drug-polymer thin films were subsequently hydrated with sterile saline and incubated at 37 °C for 10 min. Drug loadings within the micellar formulations were assessed employing a high-pressure liquid chromatography (HPLC) system (Agilent Technologies 1200 series) equipped with a Nucleosil C18 column (4.6 mm ×250 mm, 5 μm). Micelle formulations were diluted 50-fold with mobile phase, and 10 μL of diluted micelle samples were injected into the HPLC system for drug loading analysis. The mobile phase consisted of water (0.1% trifluoroacetic acid) and acetonitrile (0.1% trifluoroacetic acid) (50/ 50 vol ratio). The flow rate of the mobile phase was maintained at 1.0 mL/min. The detection wavelengths were 280 nm for Res and ID. The drug loading efficiency (LE = M _drug_ / M _drug added_ × 100) and drug loading capacity (LC = M _drug_ / (M _drug_ + M _excipient_) × 100) were calculated for each of the micelle formulations. The Zetasizer Nano ZS (Malvern Instruments Ltd., UK), equipped with a multi-angle sizing option, was employed to determine the size distribution of micelles. The effective diameter (D_eff_) and Polydispersity Index (PDI) for each micelle were ascertained by Dynamic Light Scattering (DLS) based on three measures of three independently prepared micelle drugs. The morphology of polymer micelle nanoformulation was investigated using a LEO EM910 TEM operating at 80 kV (Carl Zeiss SMT Inc., Peabody, MA). One drop of dilute sample was dropped on copper grid/carbon film and stained with 1% uranyl acetate before the TEM imaging. Digital images were obtained using a Gatan Orius SC1000 CCD Digital Camera in combination with Digital Micrograph 3.11.0 software (Gatan Inc., Pleasanton, CA). The drug release profile was assessed by the membrane dialysis method under sink conditions. The concentration of nanoformulations was adjusted to 0.1 g/L of total drug concentration in the in phosphate-buffered saline (PBS). The diluted micelle drug solutions were then moved to floatable Slide-A-Lyzer MINI dialysis devices (100 μL capacity, 3.5 kDa MWCO) and placed in 30mL of PBS complemented with 10% FBS. At predetermined time points, the remaining drugs in the micelle drug solutions were measured by HPLC, as described above.

### In vivo toxicity studies of free drugs and nanoformulated drugs

All animal procedures were conducted in accordance with the University of North Carolina at Chapel Hill Institutional Animal Care and Use Committee (IACUC) guidelines. Healthy 6-8 weeks old female BALB/c mice were obtained from Jackson Laboratory. The mice (5 mice/cage) were located in a husbandry room for two weeks prior to the study with free access to water and food under 20% humidity, 72–74°F temperature, and a 12 h light/dark cycle. The cages were randomly assigned to control and treatment groups. The dose-escalation toxicity study was conducted with freshly prepared free drug formulations (dissolved in 10% DMSO, 40% PEG300, and 5% Tween 80 in distilled water), single-loaded, or co-loaded micelles in 0.9% saline solution either through tail vein intravenous (i.v.) administration or intraperitoneal injection (i.p.) using q4d × 4 dosing regimens. The mice were injected with free drug formulations: vehicle, Res (5 mg/kg), Res (10 mg/kg), Res (20 mg/kg), ID (20 mg/kg), ID (40 mg/kg), ID (80 mg/kg), Res:ID (5:20 mg/kg), Res:ID (10:40 mg/kg), and Res:ID (20:80 mg/kg). Next, a new cohort of mice were injected with single-loaded or co-loaded micelles: Saline, Res (20 mg/kg), ID (80 mg/kg), and Res:ID (20:80 mg/kg). The general behavior and body weight changes of mice were monitored and recorded biweekly for up to 45 days. Drug treatments were discontinued if any signs of toxicity including hunched posture, rough coat, and body weight changes greater than 20% of the initial body weight. For hematology and blood chemistry studies, blood was collected into K2-EDTA microtainer (500µl) tubes (BD 365974, USA) with cardiac puncture using 25 G syringes. Cell blood count and hematology parameters were determined by IDEXX Procyte DX (Westbrook, Maine, USA). The plasma of the blood samples was obtained by centrifuging at 2,000 G for 15 min, and the blood chemistry parameters were determined using a Alfa Wassermann Vet Axcel (West Caldwell, NJ, USA) blood chemistry analyzer. For histopathological assessment, organs (Spleen, Liver, Kidney, and Lung) were harvested, and preserved in 10% neutral buffered formalin (Sigma-Aldrich), for further histological examination. The paraffin processing and embedding was performed by the Center for Gastrointestinal Biology and Disease core at UNC. Tissues were processed on the standard cycle at 60°C in a Leica ASP300S Tissue Processor for approximately 8.5 hours. After completion of overnight processing, samples were embedded using the Leica EG1160 Embedding Station. Blocks were sectioned using a Leica RM2235 Microtome by Histology Research Core Facility at UNC. 5 micron sections were floated on a water bath before being placed on charged slides. Slides were allowed to air dry overnight and were then baked at 60C for 1.5 hours. Slides were cooled overnight at room temperature prior to with Hematoxylin and Eosin staining.

### In vivo tumor inhibition studies

For tumor inoculation, T11-AP cells were mixed with a 1:1 vol ratio with Matrigel (Corning, AZ) mixture, and mice were inoculated orthotopically with 1 ×10^5^ T11-AP cells in the 2^nd^ mammary fat pad. Mice were randomized when the tumor reached 100 mm^3^ in size. For the groups receiving RT treatment, tumors were irradiated with 8 Gy photon radiation delivered using a Precision X-RAD 320 (Precision X-ray) machine operated at a peak voltage of 320 kV and at 12.5 mA. To obtain targeted RT, a lead shield protected the mice, and only the tumor was exposed to radiation. The tumors were irradiated on days 4, 5, and 6, and mice received the drug formulations by i.v. injections via tail vein using q4d × 4 regimen. Tumor size was closely monitored biweekly. Tumor length (L) and width (W) were measured by caliper, and tumor volume (V) was calculated by V = 1⁄2 x L x W^2^ equation. For free drug administration by oral gavage and i.v. injection, drugs were solubilized in a mixture of 10% DMSO, 40% PEG300, and 5% Tween 80 and distilled water. Tumor measurement continued until the end-point (tumor diameter ≥ 2cm). In depletion study, antibody was delivered bi-weekly using 100ul volumes i.p. injection for the following: aCD8 (500ug, Bioxcel BE0004), aCD4 (500ug, Bioxcel BE0003), anti-CD19 (400ug, Bioxcel BE0150), and Ly6G/Ly6C (Gr-1) (200ug, Bioxcel BE0075). Mice received one tail vein i.v. injection of aCD20 (Ultra-LEAF Purified anti-mouse CD20, 152104).

### Flow cytometry

The flow cytometry-based panels were designed and validated for comprehensive immunophenotyping of the major immune cell subsets in the tumor and tumor-draining lymph node (TDLN) of tumor-bearing mice. Orthotopic T11-AP tumors from Balb/c mice were harvested on predetermined time points and processed to obtain single-cell suspensions through mechanical and enzymatic digestion. Initially, resected tumors were divided into small pieces using a razor in a 35x10mm cell culture Petri dishes, and consequently, tissues were digested by a gently stirred digestion cocktail (collagenase, DNase I, hyaluronidase (Sigma-Aldrich), and Liberase (Roche)) for 45 minutes in an incubator at 37°C. The digested tissues were then filtered through a 70 μm cell strainer, washed with staining buffer, and resuspended in 1mL red blood cell lysis buffer (ammonium-chloride-potassium). The obtained single-cell suspensions were washed with PBS and incubated with fixable live/dead stain for 30 minutes on ice. The live/dead stained single-cell suspensions were then incubated with Fc Block (clone 2.4G2; Bio X Cell) solution on ice for 20 minutes and subsequently incubated with a cocktail of fluorochrome-conjugated antibodies for 25 minutes at 4 °C. The cell suspension was washed by staining buffer and was fixed/ permeabilized using Foxp3 Staining Buffer Set, according to the manufacturer’s instructions (ThermoFisher). Subsequently, fixed/permeabilized cells were stained with the intercellular antibodies for 30 minutes at 4 °C, washed with permeabilization buffer, and resuspended in PBS to be analyzed on flow cytometry instrument. The list of antibodies used for flow cytometry are listed in supplementary Table 1S. Reference samples were prepared using UltraComp eBeads^TM^ Compensation Beads (ThermoFisher) or tissue-specific single-cell suspensions by individually staining with fluorochrome-conjugated antibodies. The samples were analyzed using either the BD FACSymphony™ A3 Cell Analyzer equipped with FACSDiva software or 5-laser Cytek® Aurora (Cytek® Biosciences). The obtained data were analyzed by FlowJo software (TreeStar, Ashland, OR) using the gating strategy illustrated in Figure 1S. Furthermore, OMIQ software was utilized to achieve the high-dimensional analysis (https://www.omiq.ai/).

### Statistical analysis

Data are presented as mean ± SEM, as stated in the figure legends. Statistical significance was determined as indicated in the figure legends using GraphPad Prism 9 software, with P < 0.05 being considered statistically significant, *P < 0.05, **P < 0.01, ***P < 0.001, and ****P < 0.0001.

## Acknowledgments

Financial support was provided by the Basic Immune Mechanisms Training Program (NIH T32AI007273), the Carolina Cancer Nanotechnology Training Program (NIH T32CA196589), TTNCI (NIH R01 CA2644488) (all to MY), the ROI Grant from the State of North Carolina (JSS), and and the NCI Breast SPORE program (JSS) (P50-CA058223) and RO1-CA148761 (CMP). Some figures were created with BioRender.com. Animal Studies were performed within the UNC Lineberger Animal Studies Core (ASC) Facility at the University of North Carolina at Chapel Hill. The authors thank M. Ross and P. V. Kantun at the Animal Studies Core of UNC for helping with the oral gavage and intravenous injections. The UNC Flow Cytometry Core Facility is supported in part by P30 CA016086 Cancer Center Core Support Grant to the UNC Lineberger Comprehensive Cancer Center. The authors thank Dr. Ramiro Diz at UNC Flow Cytometry Core Facility for his technical assistance and critical feedback. TEM was performed by A. Shankar Kumbhar at the Chapel Hill Analytical and Nanofabrication Laboratory (CHANL), a member of the North Carolina Research Triangle Nanotechnology Network (RTNN), which was supported by the NSF (grant ECCS-1542015) as part of the National Nanotechnology Coordinated Infrastructure (NNCI). Histological services provided by the Histology Research Core Facility in the Department of Cell Biology and Physiology at the University of North Carolina, Chapel Hill NC. We acknowledge Dr. Rani Sellers at Department of Pathology and Laboratory Medicine of the University of North Carolina, Chapel Hill NC for histological analysis.

## Authorship contribution statement

Mostafa Yazdimamaghani: Conceptualization, Data curation, Methodology, Writing, Visualization.

Oleg Kolupaev: Conceptualization, Data curation, Methodology, Writing – review & editing, Visualization. Chaemin Lim: Methodology, Visualization, Writing – review & editing.

Duhyeong Hwang: Methodology, Validation.

Sonia J. Laurie: Methodology, Writing – review & editing.

Charles M. Perou: Writing – review & editing, Resources.

Alexander V. Kabanov: Writing – review & editing, Resources.

Jonathan S. Serody: Conceptualization, Writing – review & editing, Project administration, Funding acquisition, Resources.

## Data Availability Statement

The data that support the findings of this study are available from the corresponding author upon reasonable request.

