## Supplementary Material for "Tumor microenvironment immunomodulation by nanoformulated TLR 7/8 agonist and PI3k delta inhibitor enhances therapeutic benefits of radiotherapy"

| Antibody | Manufacturer | Clone | Catalog number |
|---|---|---|---|
| LIVE/DEAD™ Fixable Blue Dead Cell Stain Kit | Invitrogen | | L23105 |
| FITC anti-mouse TER-119/Erythroid Cells | BioLegend | TER119 | 116205 |
| FITC anti-mouse/human CD11b | BioLegend | M1/70 | 101206 |
| NK1.1 Monoclonal Antibody, FITC | eBioscience | PK136 | 11-5941-82 |
| Ly-6G/Ly-6C Monoclonal Antibody, FITC | eBioscience | RB6-8C5 | 11-5931-82 |
| FITC anti-mouse CD11c | BioLegend | N418 | 117305 |
| Brilliant Violet 510™ anti-mouse CD45 | BioLegend | 30-F11 | 103137 |
| BUV395 Hamster Anti-Mouse TCR β Chain | BD Biosciences | H57-597 | 742485 |
| BUV737 Rat Anti-Mouse CD4 | BD Biosciences | RM4-5 | 612844 |
| APC/Cyanine7 anti-mouse CD8a | BioLegend | 53-6.7 | 100714 |
| CD62L (L-Selectin) Monoclonal Antibody, PE-eFluor™ 610 | eBioscience | MEL-14 | 61-0621-82 |
| Brilliant Violet 711™ anti-mouse/human CD44 | BioLegend | IM7 | 103057 |
| Brilliant Violet 785™ anti-mouse CD103 | BioLegend | 2E7 | 121439 |
| BB700 Rat Anti-Mouse CD25 | BD Biosciences | PC61 | 566499 |
| Brilliant Violet 421™ anti-mouse CD279 (PD-1) | BioLegend | 29F.1A12 | 135221 |
| APC anti-mouse CD366 (Tim-3) | BioLegend | B8.2C12 | 134007 |
| Alexa Fluor® 700 anti-mouse CD19 | BioLegend | 6D5 | 115528 |
| PE/Cyanine7 anti-mouse IgM | BioLegend | RMM-1 | 406514 |
| BUV805 Mouse Anti-Mouse IgD | BD Biosciences | AMS 9.1 | 748469 |
| BV605 Rat Anti-Mouse IgG1 | BD Biosciences | X56 | 742477 |
| FOXP3 Monoclonal Antibody, PE | eBioscience | FJK-16S | 12-5773-82 |
| Brilliant Violet 650™ anti-mouse I-A/I-E | BioLegend | M5/114.15.2 | 107641 |
| Brilliant Violet 605™ anti-mouse CD8a | BioLegend | 53-6.7 | 100743 |
| Spark UV™ 387 anti-mouse/human CD11b | BioLegend | M1/70 | 101291 |
| Brilliant Violet 785™ anti-mouse Ly-6C | BioLegend | HK1.4 | 128041 |
| BUV805 Rat Anti-Mouse Ly-6G | BD Biosciences | 1A8 | 741994 |
| PE/Cyanine7 anti-mouse F4/80 | BioLegend | BM8 | 123113 |
| BB700 Rat Anti-Mouse CD86 | BD Biosciences | GL1 | 742120 |
| BUV563 Rat Anti-Mouse CD45R/B220 | BD Biosciences | RA3-6B2 | 748868 |
| PE/Dazzle(TM) 594 anti-mouse CD163 | BioLegend | S15049F | 156709 |
| PE/Cyanine5 anti-mouse CD103 | BioLegend | 2E7 | 121447 |
| iNOS Monoclonal Antibody, Brilliant Violet™ 421 | eBioscience | CXNFT | 404-5920-82 |
| Alexa Fluor® 647 Rat Anti-Mouse CD206 | BD Biosciences | MR5D3 | 565250 |
| InVivoMAb anti-mouse CD19 | Bioxcell | 1D3 | BE0150 |
| InVivoMAb anti-mouse Ly6G/Ly6C (Gr-1) | Bioxcell | RB6-8C5 | BE0075 |
| InVivoMAb anti-mouse CD4 | Bioxcell | GK1.5 | BE0003-1 |
| InVivoMAb anti-mouse CD8α | Bioxcell | 53-6.7 | BE0004-1 |
| Ultra-LEAF™ Purified anti-mouse CD20 | BioLegend | SA271G2 | 152116 |
| Brilliant Violet 510™ anti-mouse CD8a | BioLegend | 53-6.7 | 100751 |
| BUV563 Rat Anti-Mouse TER-119/Erythroid Cells | BD Biosciences | TER-119 | 741257 |
| BUV661 Rat Anti-CD11b | BD Biosciences | M1/70 | 612977 |
| BUV661 Rat Anti-Mouse IgM | BD Biosciences | II/41 | 750660 |
| BUV737 Rat Anti-Mouse CD4 | BD Biosciences | GK1.5 | 612761 |
| Alexa Fluor® 647 anti-mouse CD185 (CXCR5) | BioLegend | L138D7 | 145532 |
| BB700 Rat Anti-Mouse CD45R (B220) | BD Biosciences | RA3-6B2 | 746206 |
| APC/Fire™ 810 anti-mouse Ly-6C | BioLegend | HK1.4 | 128055 |
| PE anti-mouse CD84 | BioLegend | mCD84.7 | 122805 |
| BV750 Rat Anti-Mouse CD138 | BD Biosciences | 281-2 | 747070 |
| Brilliant Violet 785™ anti-mouse CD25 | BioLegend | PC61 | 102051 |
| PerCP anti-mouse CD45 | BioLegend | 30-F11 | 103129 |
| FOXP3 Monoclonal Antibody, APC | eBioscience | FJK-16s | 17-5773-82 |
| PE/Dazzle™ 594 anti-mouse CD62L | BioLegend | MEL-14 | 104447 |
| PE/Cyanine5 anti-mouse CD19 | BioLegend | 1D3 | 115509 |
| iNOS Monoclonal Antibody, PE-Cyanine7 | eBioscience | CXNFT | 25-5920-80 |
| Spark NIR™ 685 anti-mouse IgD | BioLegend | 11-26c.2a | 405749 |
| CD101 Monoclonal Antibody, Alexa Fluor™ 700 | eBioscience | Moushi101 | 56-1011-82 |
| APC/Cyanine7 anti-mouse CD103 | BioLegend | 2 E7 | 121431 |
| BUV395 Rat Anti-Mouse CD3 molecular complex | BD Biosciences | 17A2 | 740268 |

**Supplementary Table 1.** List of antibodies used in flow cytometry studies.

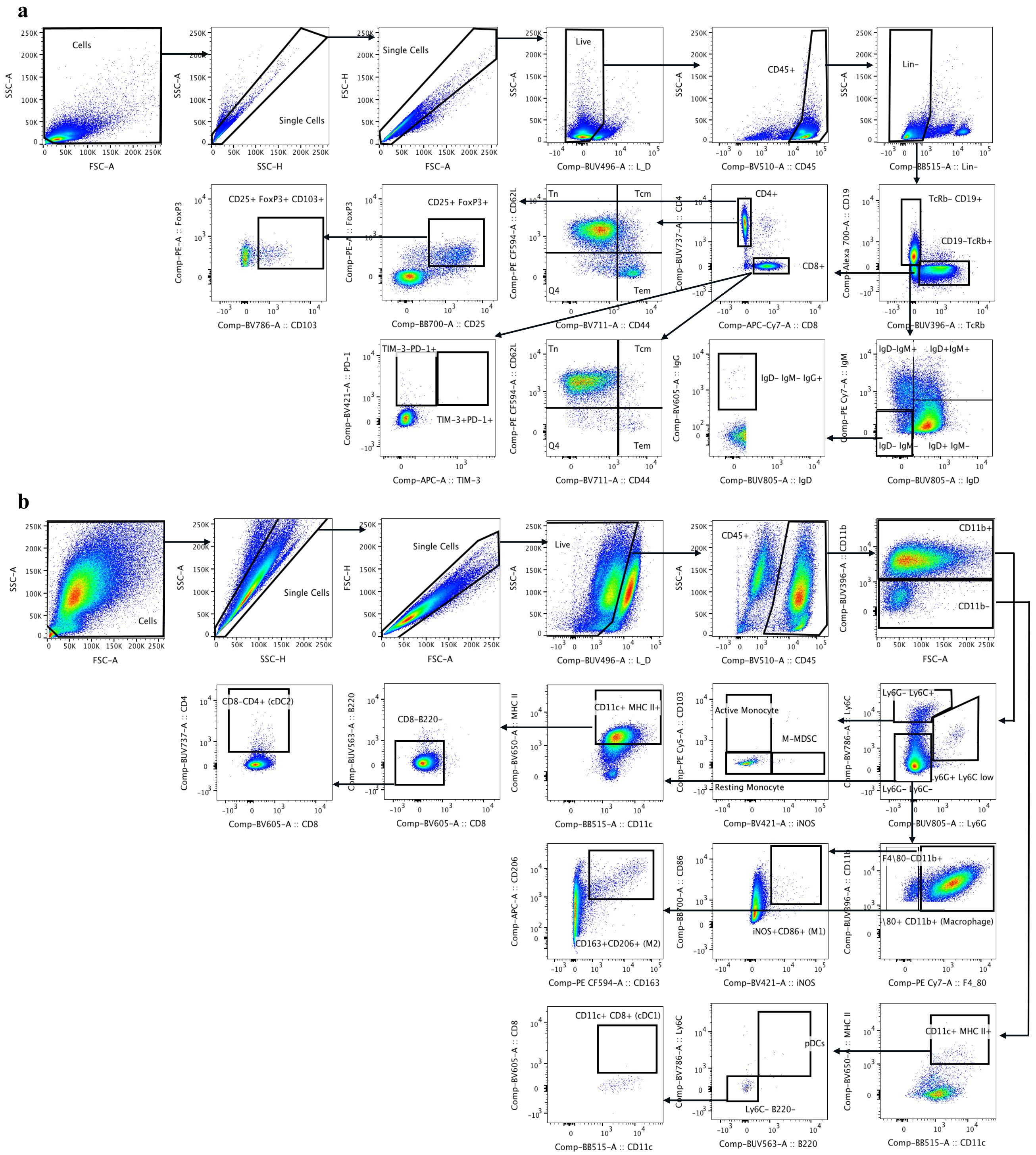


**Supplementary Figure 1. Flow Cytometry Gating Strategy a)** Gating strategy for lymphocytes isolated from the T11-AP model to examine B cells, Tregs, and T cells exhaustion state and memory phenotype. **b)** Gating strategy for myeloid cells isolated from the T11-AP model to profile M-MDSC, cDC1, cDC2, pDCs, and macrophage polarization. Fluorescence Minus One (FMO) control was used to assign gating properly.

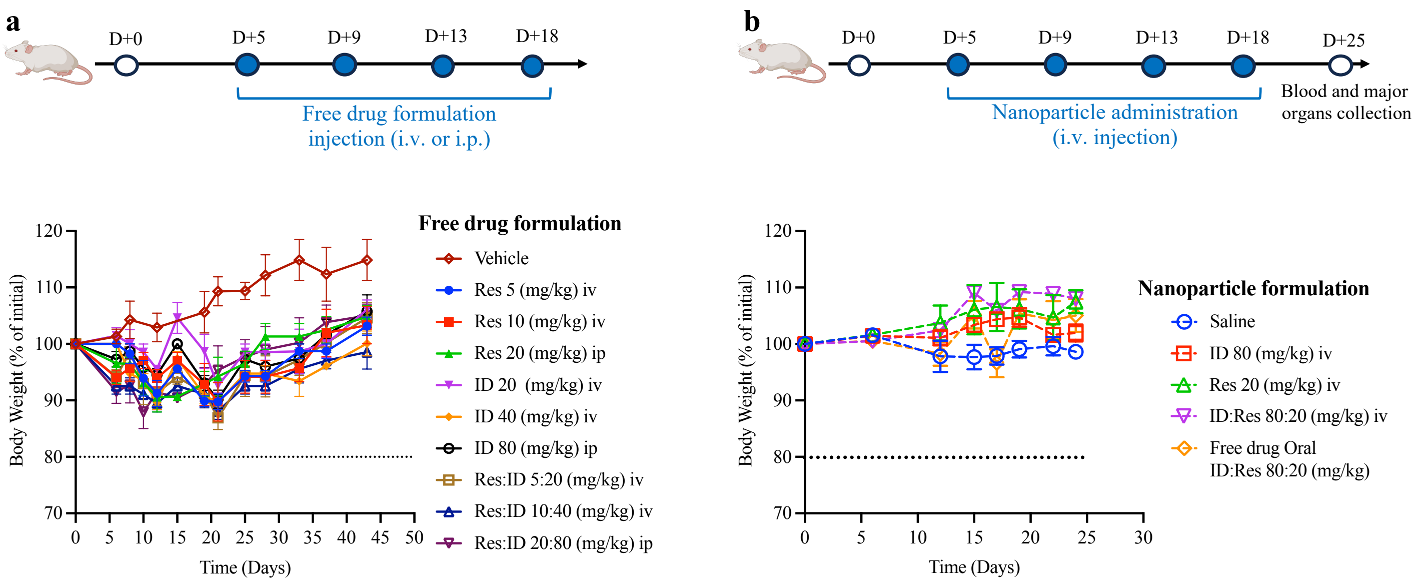


**Supplementary Figure 2. Toxicity studies of free drugs and nanoformulated drugs in healthy non-tumor-bearing mice. a)** Treatment schedule and body weight change of healthy mice injected i.v. or i.p. with free drug formulations (dissolved in 10% DMSO, 40% PEG300, and 5% Tween 80 in distilled water) (n=3). **b)** Treatment schedule and body weight change of healthy mice injected with nanoformulated POx-Res (20 mg/kg), POx-ID (80 mg/kg), POx-Res:ID (20 mg/kg Res:80 mg/kg ID), and oral delivery of free drug formulation of Res:ID (20 mg/kg Res:80 mg/kg ID) (n=3).

a

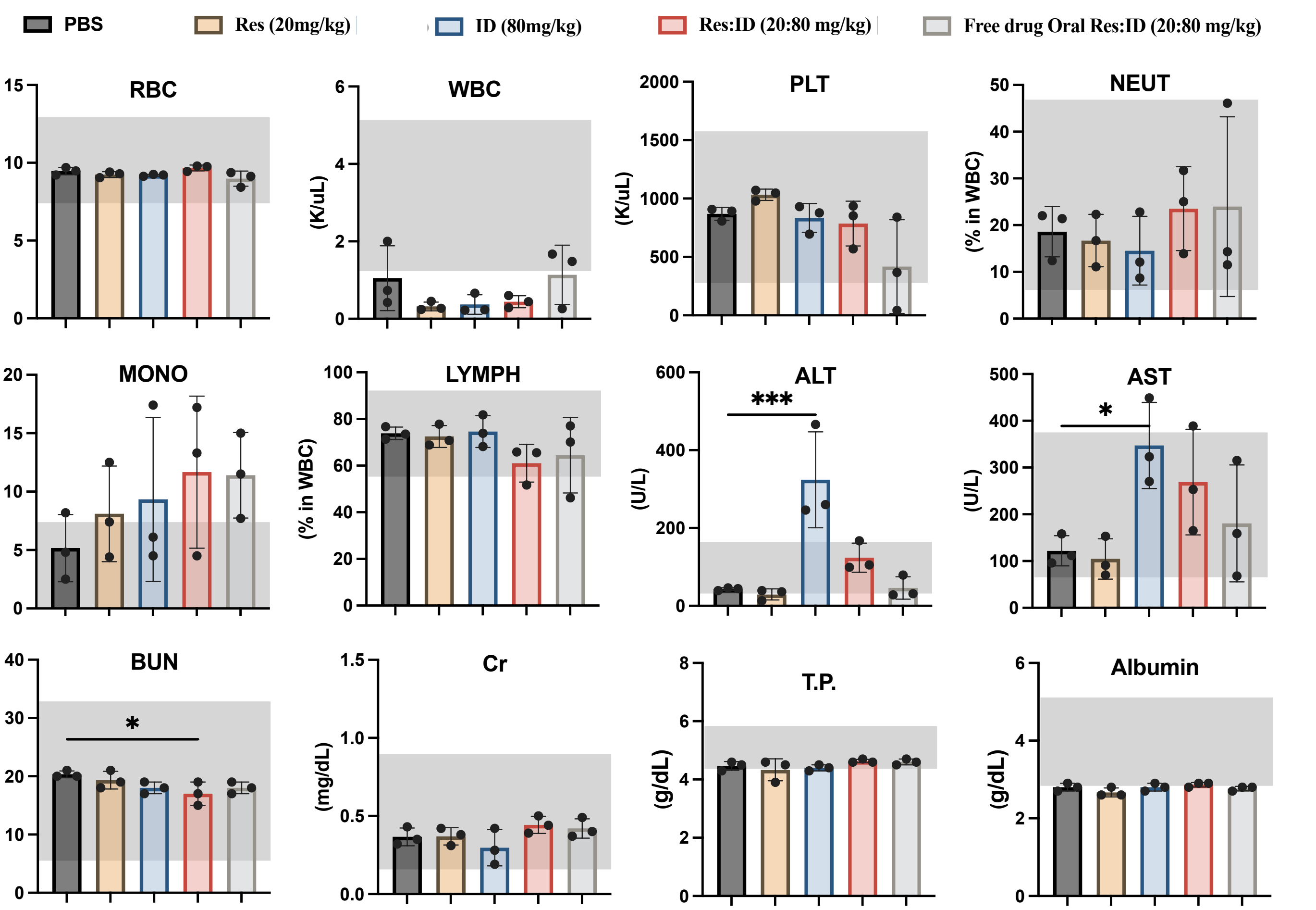


| | | Saline | Res(20mg/kg) | ID (80mg/kg) | Res:ID (20:80 mg/kg) | Free drug Oral Res:ID (20:80 mg/kg) | Normal range |
|---|---|---|---|---|---|---|---|
| Complete blood count | RBC (M/uL) | 9.5±0.2 | 9.2±0.2 | 9.2±0.1 | 9.7±0.2 | 9±0.5 | 7~12 |
| | WBC (K/uL) | 1.1±0.8 | 0.3±0.1 | 0.4±0.3 | 0.4±0.2 | 1.1±0.8 | 1.8~5.2 |
| | PLT (K/uL) | 869.3±54.5 | 1032.3±47.7 | 834±123.3 | 785.3±191.9 | 417±401.7 | 285~1543 |
| | Neutro (% in WBC) | 18.6±5.4 | 16.7±5.6 | 14.5±7.4 | 23.5±9 | 24±19.2 | 8~48 |
| | Mon (% in WBC) | 5.2±2.9 | 8.1±4.1 | 9.3±7 | 11.7±6.5 | 11.4±3.7 | 0~7 |
| | Lym (% in WBC) | 73.9±2.7 | 72.5±4.7 | 74.6±6.8 | 61±8.1 | 64.4±16.2 | 57~93 |
| Blood chemistry | ALT (U/L) | 43±3.6 | 29.3±14.2 | 324.3±123.2 *** | 123.7±37.6 | 46±28.6 | 40-170 |
| | AST (U/L) | 122±32.2 | 104.7±43.2 | 347.3±91.9 * | 269±112.9 | 180.7±124.9 | 67-381 |
| | BUN (mg/dL) | 20.3±0.6 | 19.3±1.5 | 18±1 | 17±2 * | 18±1 | 7-31 |
| | Cr (mg/dL) | 0.4±0.1 | 0.4±0.1 | 0.3±0.1 | 0.4±0.1 | 0.4±0.1 | 0.2~0.9 |
| | T.P (g/dL) | 4.5±0.2 | 4.3±0.4 | 4.4±0.1 | 4.6±0.1 | 4.6±0.1 | 4.4~5.8 |
| | Albumin (g/dL) | 2.8±0.1 | 2.7±0.1 | 2.8±0.1 | 2.9±0.1 | 2.8±0.1 | 2.7~5.3 |

b

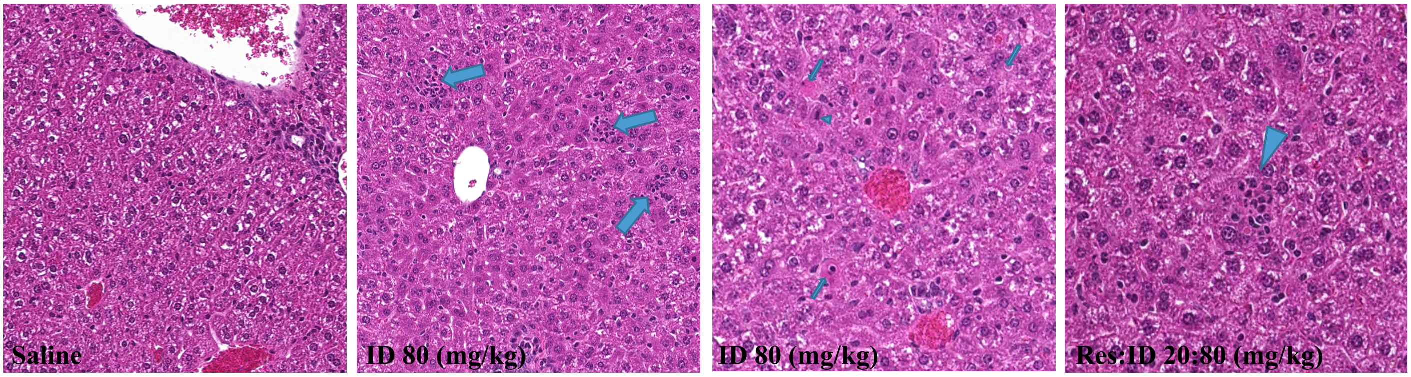

**Fig. 3S. Hematological, biochemical, and histological effects of Oral free drugs and nanoformulated drugs in healthy non-tumor-bearing mice. a)** Complete blood count and clinical blood chemistry parameters of healthy mice treated with saline and POx-Res (20 mg/kg), POx-ID (80 mg/kg), POx-Res:ID (20 mg/kg Res:80 mg/kg ID), and oral delivery of free drug formulation of Res:ID (20 mg/kg Res:80 mg/kg ID) (n=3) were assayed at 7 days after the last injection. RBC, red blood cells ($10^{12}$/liter); WBC, white blood cells ($10^{9}$/liter); PLT, platelets ($10^{9}$/liter); Neutro, neutrophils (% in WBC); Mon, monocytes (% in WBC); Lym, lymphocytes (% in WBC); ALT, alanine aminotransferase (U/liter); AST, aspartate aminotransferase (U/liter); T.P., total protein (g/dl); BUN, blood urea nitrogen (mg/dl); Cr, creatinine (mg/dl). Data are mean ± SD (n = 3). All data are presented as the mean ± SD. Ordinary one-way ANOVA followed by Dunnett's multiple-comparisons test was applied for statistical analyses. *P*values: NS, not significant, $*P < 0.05$, $**P < 0.01$, $***P < 0.001$, $****P < 0.0001$). **b)** Histological analysis of major organs (Spleen, Liver, Kidney, and Lung). One week after the last injection, major organs were harvested for H&E and Masson's trichrome staining and histological analysis. A licensed pathologist evaluated histological changes.

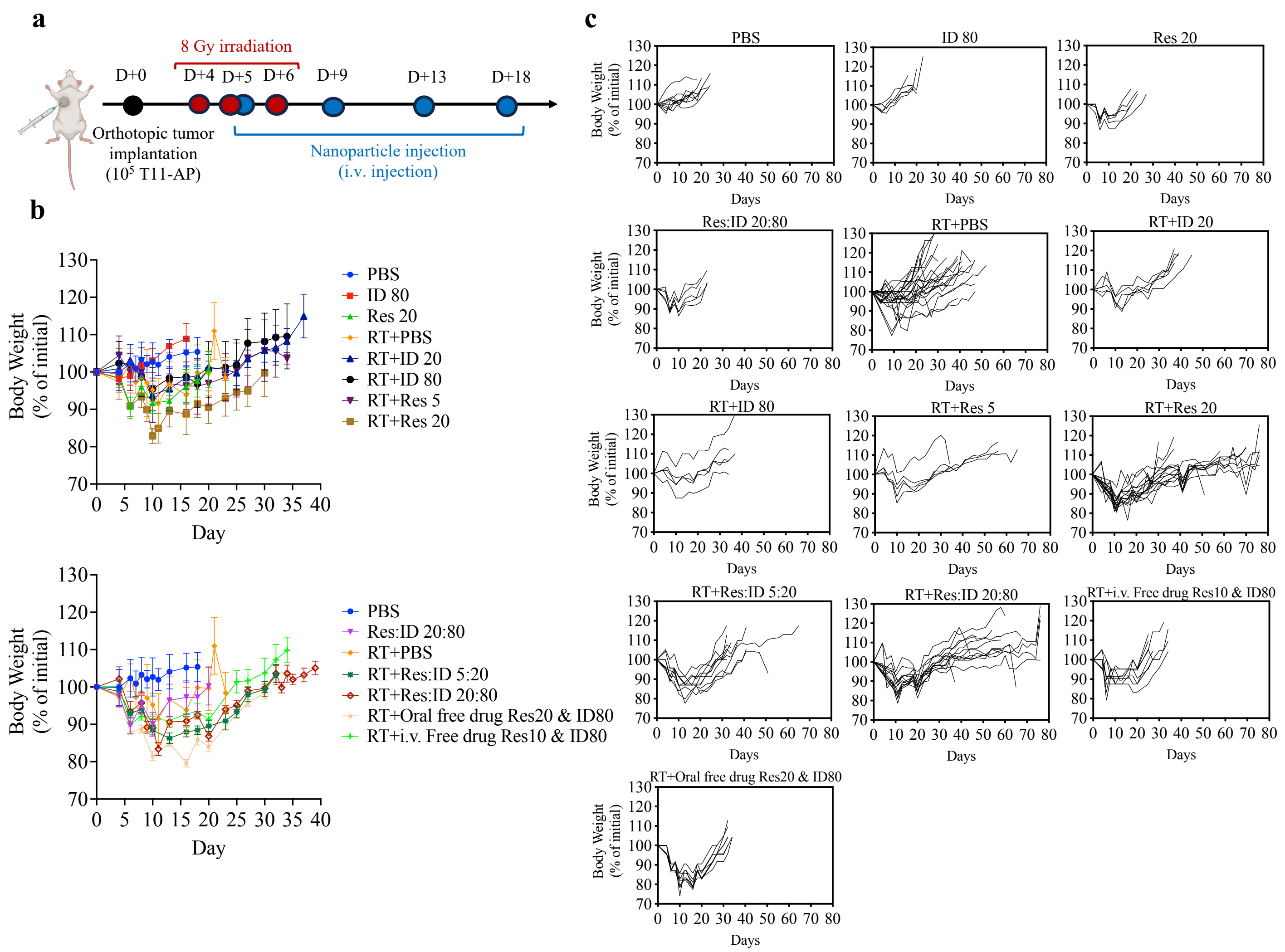


**Supplementary Figure 4. Body weight changes over the course of treatment. a)** Treatment schedule of tumor-bearing mice. **b)** Average (mean ± SEM) body weight changes and **c)** individual percentage of body weight changes for different treatment groups.

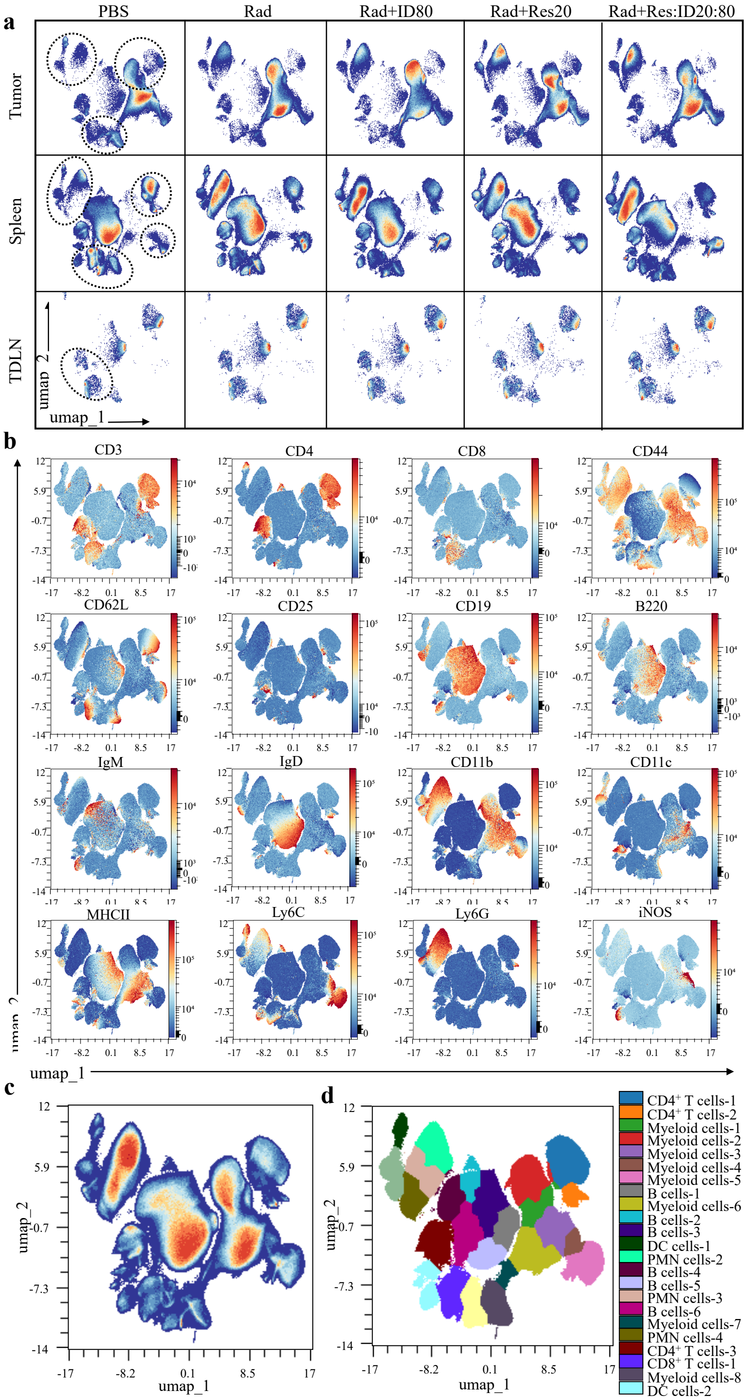


**Supplementary Figure 5. High dimensional spectral flow cytometry analysis for an in-depth investigation of immune landscape profile across different treatments in tumor and TDLN.** Each dot represents a single cell. In total, $4.3 \times 10^6$ $CD45^{+}TER119^{-}$ live immune cells were analyzed by OMIQ**. a)** Representative UMAP plots of tumor and TDLN tissues, comparing control, RT, combined RT with ID80 NPs, combined RT with Res20 NPs, and combined RT with Res:ID20:80 NPs subjected to 26-parameter phenotyping. **b)** Relative canonical marker expression of major lineages. **c)** Representative UMAP density plot and **d)** FlowSOM Clusters projected on UMAP plot of all identified myeloid and lymphoid cells.

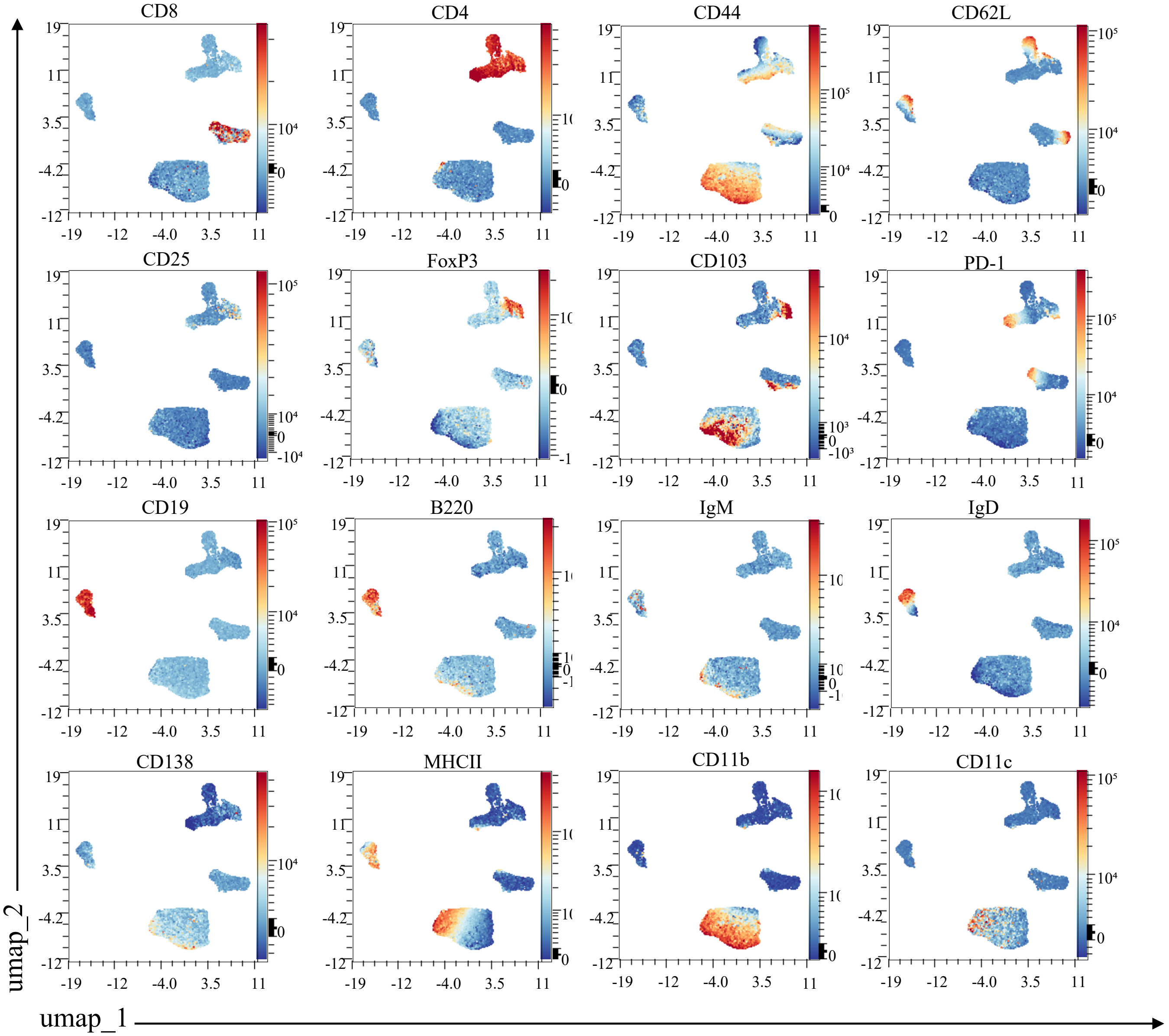


**Supplementary Figure 6. Representative UMAP scatterplot of relative expression of major lineages markers of tumor lymphocytes.**

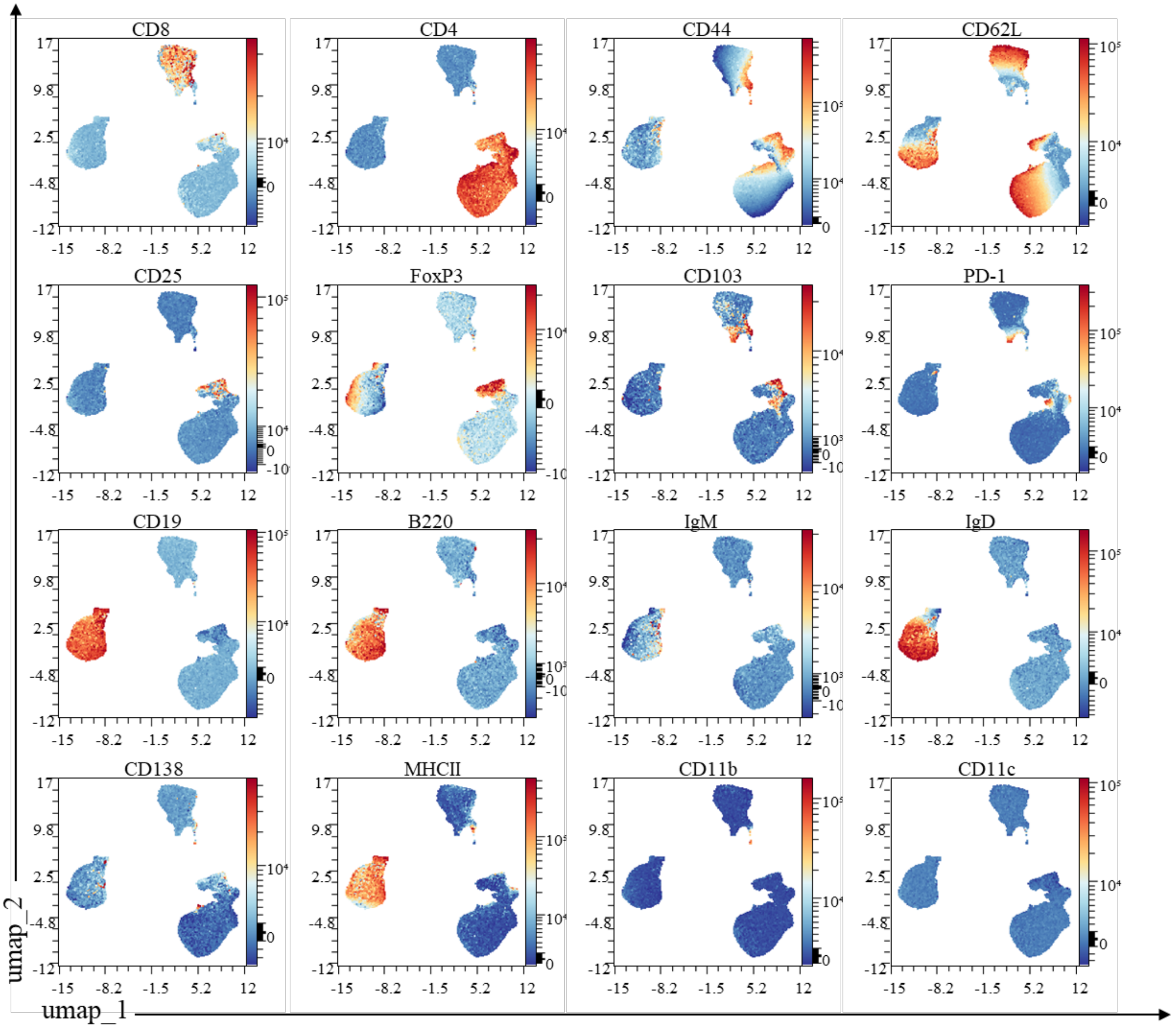


**Supplementary Figure 7. Representative UMAP scatterplot of relative expression of major lineages markers of TDLN lymphocytes.**

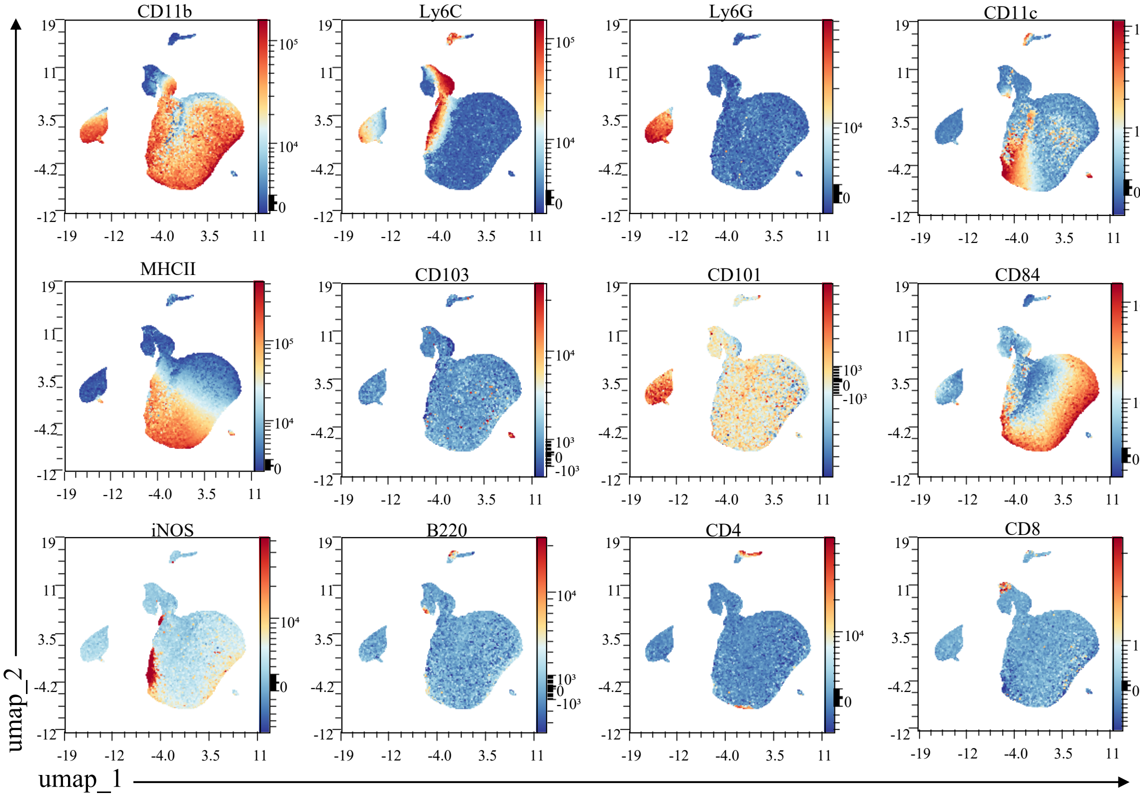


**Supplementary Figure 8. Representative UMAP scatterplot of relative expression of major lineages markers of tumor myeloid cells.**
